# Systematic identification of cell division proteins in haloarchaea and the discovery of a membrane anchor for the Z ring

**DOI:** 10.64898/2026.07.16.738847

**Authors:** Wenchao Zheng, Shan Zhao, Yafei Liu, Jiayao Cui, Jiarong Wu, Xiaoyao Zou, Ying Li, Han Gong, Xiangdong Chen, Joe Lutkenhaus, Fabai Wu, Shishen Du

## Abstract

Most archaea rely on the tubulin-like protein FtsZ for division. In the last decade, several novel cell division proteins have been discovered in the model haloarchaeon *Haloferax volcanii* which contains two FtsZ proteins, FtsZ1 and FtsZ2. However, the composition of this FtsZ-based archaeal divisome is largely uncharacterized. Here, using *in vivo* crosslinking coupled with immunoprecipitation and mass spectrometry (CLIP-MS), we identified ten proteins that localize to the division site, including wide-spread proteins predicted to be involved in DNA binding and energy metabolism. Deletion analysis indicate that most of these proteins have a modest or minor impact on cell division, but HVO_0399, renamed as <u>C</u>ell <u>D</u>ivision <u>P</u>rotein <u>C</u> (CdpC), is important for cell division and functions as a membrane anchor for FtsZ1. CdpC consists of three domains, an N-terminal domain (NTD) containing an amphipathic helix critical for membrane binding, a long intrinsically disordered linker, and a C-terminal domain (CTD) that binds FtsZ1 with high affinity and likely promotes its polymerization. Notably, both the NTD and CTD of CdpC harbor a zinc finger that is indispensable for its function. Phylogenetic analysis indicates that CdpC sequences evolved fast across haloarchaea, but exhibited high conservation in domain structure and critical residues found in this study. Overall, our study expands the repertoire of candidate division proteins and establishes CdpC as a membrane anchor for FtsZ1 in haloarchaea. These findings pave the way for in-depth studies of arcaheal cell division and broaden the function of zinc finger proteins in archaea.

## Main

Cell division is one of the most fundamental biological processes for all cellular organisms. Across the tree of life, different life forms rely on distinct mechanisms for division. Most bacteria employ the tubulin-like protein FtsZ for division, while archaea can divide by the FtsZ-based system or the Endosomal Sorting Complex Required for Transport III-based (ESCRT-III) system^1–10^.^9^[ref9] Evolutionary analysis of FtsZ revealed that it is widely present in prokaryotes and dates back to the Last Universal Common Ancestor (LUCA), indicating that the FtsZ-based system may be the archetypical ancestral division apparatus^8,11^. Interestingly, while bacteria typically encode a single FtsZ protein, many archaea possess two FtsZ paralogs, FtsZ1 and FtsZ2^8^. FtsZ is a GTPase which assembles into a ring-like structure, the Z ring, with the aid of its membrane anchors and associated proteins, at midcell to initiate cytokinesis in both bacteria and archaea^8,12–14^. Once this cytoskeletal element is formed, it functions as a scaffold for the assembly of the division apparatus (or the divisome) which executes cytokinesis.

The haloarchaeon *Haloferax volcanii* in Euryarchaeota, which utilizes two FtsZ proteins for division, serves as a model for the investigation of the FtsZ-based division mechanism in archaea^8,15^. Both FtsZ1 and FtsZ2 are critical for *H. volcanii* division, but their functions are distinct: FtsZ1 provides a scaffold for the divisome by assembling into the Z ring, while FtsZ2 is required for constriction^16^. In addition to FtsZ1 and FtsZ2, several cell division proteins have been identified and characterized in *H. volcanii*, including SepF, Cell Division Protein A (CdpA) and CdpB1/2/3^11,17–20^. SepF is a membrane anchor for FtsZ2 and also plays important roles in recruiting downstream proteins to the Z ring^11,17^. CdpA is a transmembrane protein organizing the divisome complex, presumably by interacting with multiple divisome proteins, especially FtsZ2 and SepF^18^. CdpB1/2/3 are small proteins belonging to the widespread PRC-barrel (Photosynthesis Reaction Center) protein family^19,20^. They interact with each other to form a complex and among them, CdpB1 directly interacts with SepF. The CdpB1-3 complex is thought to aid the organization of the Z ring and the recruitment of downstream division proteins^19,20^. The discovery and characterization of these proteins greatly increased our understanding of the composition, assembly pathway and regulation of the FtsZ-based archaeal division machine. However, many fundamental questions remained untouched. For example, how is FtsZ1 anchored to the membrane to form the Z ring in archaea relying on two FtsZs? Also, the number of identified division proteins remains limited in comparison to bacteria, hindering in-depth investigation of this FtsZ-based division mechanism in archaea.

In this study, we used a combination of biochemical and cellular approaches to systematically identify novel cell division proteins in *H. volcanii*. We successfully discovered ten divisome-associated proteins, one of which is a zinc finger protein that functions as a membrane anchor for FtsZ1 and is critical for cell division. Thus, this study significantly increases the repertoire of candidate division proteins and provides novel insights into Z ring formation in archaea.

## Results

### Systematic identification of novel cell division proteins in *H. volcanii*

Attempts to identify novel division proteins by co-immunoprecipitation (Co-IP) using a known division protein as a bait has had limited success^17^, since the divisome complex is likely highly dynamic due to the treadmilling behavior of FtsZ filaments and transient interactions among divisome components^18^. We posited that treatment of the cells with a crosslinker should stabilize the complex and facilitate subsequent isolation by Co-IP (Fig. 1a). To test this, we treated *H. volcanii* cells co-expressing 6×His tagged CdpB1 (CdpB1-6×His) and GFP tagged FtsZ1 (FtsZ1-GFP), which do not interact with each other directly^19,20^, with the crosslinker formaldehyde or 3,3’-Dithiobis(succinimidyl propionate) (DSP). After isolation of the immunocomplexes with an anti-His antibody, we checked if the crosslinked products could be detected by anti-GFP antibody. As shown in Extended Data Fig. 1, without the treatment, some faint high molecular weight bands were detected, presumably due to non-specific proteins reacting with the anti-GFP antibody. However, after treatment with formaldehyde or DSP, a high molecular weight band appeared. To ensure FtsZ1-GFP was within the immunocomplexes, we treated the samples with 1,4-Dithiothreitol (DTT) which can uncrosslink the complexes. As expected, a band corresponding to the molecular weight of FtsZ1-GFP was detected and its amount in the samples treated with DSP was substantially greater than that with formaldehyde (Extended Data Fig. 1). This result indicates that: 1) crosslinker treatment can greatly facilitate the isolation of indirect interaction partners of a division protein and 2) DSP appears more effective than formaldehyde. Therefore, we decided to use DSP for *in vivo* crosslinking followed by IP and mass spectrometry (CLIP-MS) to identify novel cell division proteins (Fig. 1a). We envisioned that such an approach can be done iteratively, and combined with further experimental validation, may lead to identification of all divisome components.

**Fig. 1.**
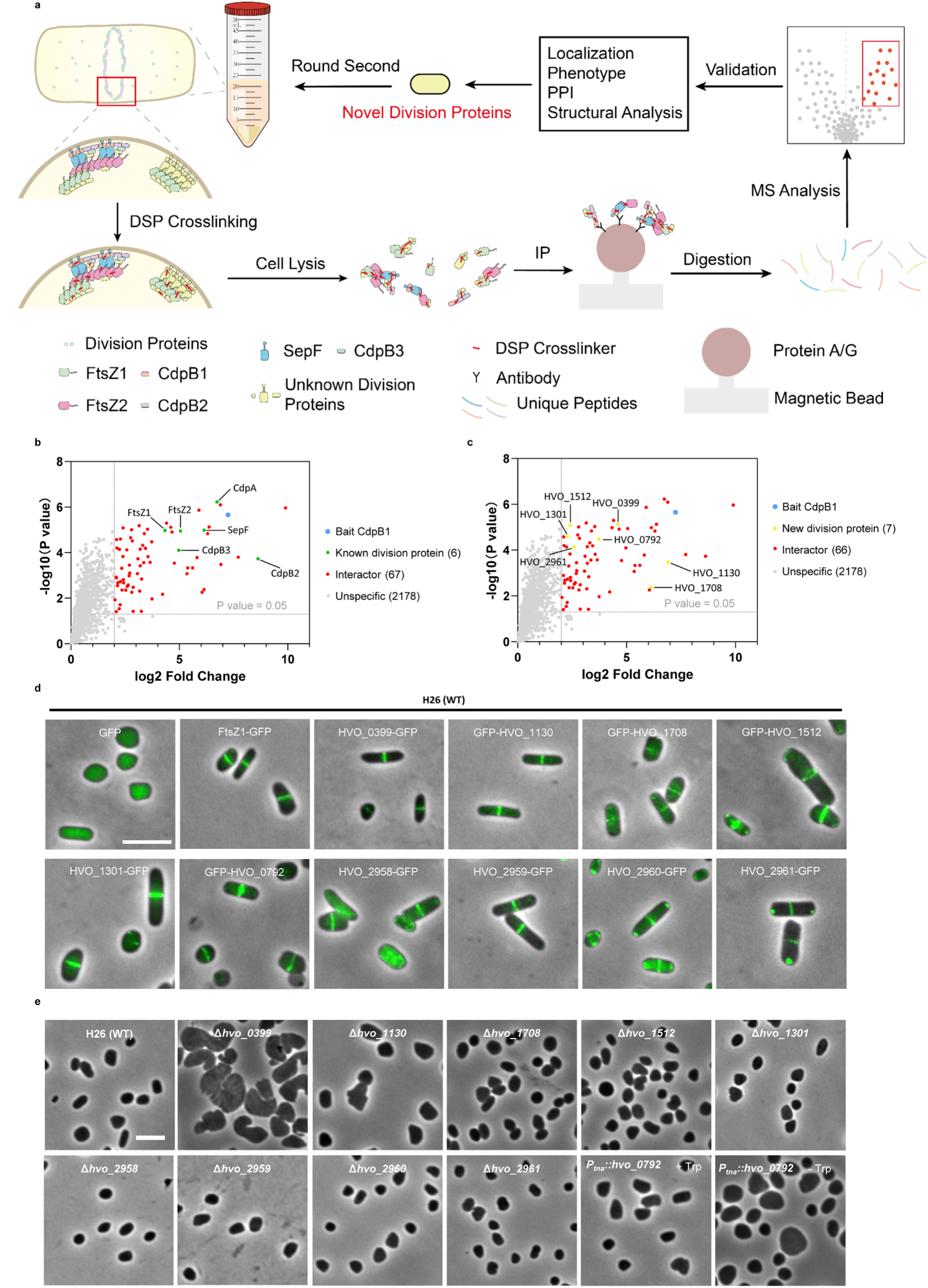
Screening for cell division proteins by *in vivo* crosslinking and IP-MS using CdpB1 as a bait in *H. volcanii*. a. Flow chart of the procedure for identification of cell division proteins in *H. volcanii* by *in vivo* cross-linking followed by IP-MS. Cells expressing CdpB1-GFP were treated with DSP to stabilize protein complexes involved in cell division. The crosslinked divisome components were enriched using anti-GFP antibody coated magnetic beads, and the co-precipitated proteins were identified by MS. Candidate proteins were further evaluated based on their subcellular localization and many other tests. Following characterization of the newly identified division proteins, additional rounds of screening can be performed to isolate further interacting partners. Arrows indicate the sequential steps of the workflow. DSP, 3,3’-Dithiobis(succinimidyl propionate); MS, mass spectrometry; PPI, protein-protein interaction. b-c. Volcano plot displays known division proteins and candidate interactors of CdpB1. Proteins with an MS enrichment ≥4-fold over the control and a P-value < 0.05 were considered as candidate interactors. All 6 known division proteins were detected (b), and 7 newly identified midcell-localized proteins were indicated in (c), along with 67 additional interactors. d. Representative images of localization of GFP fusions of ten putative new division proteins. All GFP fusion proteins were expressed from plasmids under the control of the P_tna_ promoter, induced overnight with 0.2 mM tryptophan (Trp) at 45 °C. e. Representative images of the morphology of the deletion/depletion strains of the putative new division genes. Deletion of *hvo_0399* (*cdpC*) results in severe division and morphological defects. Depletion of HVO_0792 was achieved by removal of tryptophan (Trp) from the cultures. Scale bars in (d) and (e), 5 μm.

We performed the experiments in wild type *H. volcanii* cells (H26) expressing a GFP tagged CdpB1 (Cdp1-GFP), with cells expressing GFP as a control (Fig. 1a). After isolation of the immunocomplexes, the samples were analyzed by MS. Proteins co-immunoprecipitated with CdpB1-GFP with a ≥4-fold enrichment over the GFP control and a P-value < 0.05 were considered as potential CdpB1 interaction partners, yielding 73 candidates (Supplementary Table 1). Strikingly, all the previously known division proteins—FtsZ1, FtsZ2, SepF, CdpA, CdpB2 and CdpB3—were among the hits (Fig. 1b). This indicates that CLIP-MS is highly effective in the isolation of the divisome complex or subcomplexes, with many of the enriched proteins likely to be divisome components.

To test if the 73 interactors of CdpB1 are involved in cell division, we constructed GFP fusions and examined their localization in wild type *H. volcanii* cells because localization to the midcell is a convenient visual hallmark of division proteins. Notably, seven proteins—HVO_0399, HVO_0792, HVO_1130, HVO_1301, HVO_1512, HVO_1708 and HVO_2961—displayed midcell localization (Fig. 1d). Among them, HVO_0399, HVO_0792, HVO_1130 and HVO_1131 localized almost exclusively to the midcell, while HVO_1512, HVO_1708 and HVO_2961 also exhibited additional polar foci and/or cytoplasmic fluorescence. The other proteins either displayed a uniformly distribution in the cytoplasm or formed punctates (Supplementary Table 2). As a result, they were not further characterized in the current study.

Among the seven midcell-localized proteins, HVO_2961 had been characterized as the Dihydrolipoamide dehydrogenase, the E3 component of the 2-oxoacid dehydrogenase complex (OADHC) that bridges glycolysis and the TCA cycle by converting pyruvate into acetyl-coA^21,22^. Moreover, *hvo_2961* is encoded in an operon with three other genes - *hvo_2958*, *hvo_2959*, *hvo_2960* – that encode the E1α, E1β, and E2 components of the OADHC^21^. Hence, we also examined the localization of these three proteins. Interestingly, all of them localized to midcell, with HVO_2958 exhibiting additional cytoplasmic fluorescence and HVO_2960 forming additional polar foci (Fig. 1d). Since the remaining six proteins had not been characterized previously, we predicted their functions using a sensitive hidden Markov model (HMM)-based HHpred analysis^23^ (Supplementary table 3). HVO_0399 was predicted to encode a small CPXCG-related zinc finger protein belonging to the DUF7093 family, while HVO_0792 was annotated as a 3-dehydroquinate synthase involved in the canonical shikimic pathway of aromatic amino acid biosynthesis. HVO_1130 belongs to the DUF7524 family, whereas HVO_1512 is a member of the DUF5779 family which shows similarity to SepF. HVO_1708 was predicted to be a lipid A-modifying glycosyltransferase, and HVO_1301 contains a HerA helicase domain suggesting that it may bind to nucleic acids. In summary, the *in vivo* CLIP-MS screen led to the identification of ten proteins with midcell localization (Fig. 1c), including many metabolic enzymes and a protein with putative nucleic acid binding ability.

### The absence of individual midcell-localized proteins results in cell division and shape defects to various extents

To determine if the ten midcell-localized proteins are truly involved in cell division, we generated deletion or depletion strains by the pop-in/pop-out approach ^24^. We obtained clean deletion strains for 9 genes except for *hvo_0792*, for which its promoter was replaced with the tryptophan-inducible promoter P_tna_^25^ so that its expression was regulated by tryptophan. Examination of cell morphology of the deletion strains showed that deletion of *hvo_0399* resulted in severe cell enlargement, morphological defects, and division abnormalities, while deletion of the other 8 genes had only minor effects on cell division and morphology (Fig. 1e). Depletion of HVO_0792 caused modest cell enlargement, indicating that it may be important for cell division. Taken together, these results indicate that HVO_0399 is important for division while the other midcell-localized proteins may participate in cell division, but further investigation is required to determine their role in this process.

### CdpC is important for proper Z ring formation and cell division

Given the severe phenotype of the *hvo_0399* deletion strain, we decided to focus on it in the present study. Complementation of the deletion strain with either HVO_0399 or its GFP fusion on a plasmid almost completely rescued the division and shape defects (Fig. 2a), indicating that the defects were solely caused by its absence. Moreover, HVO_0399-GFP co-localized well with mCherry fusions to FtsZ1 and FtsZ2 in wild type H26 cells (Extended Data Fig. 2a), indicating that it localizes to the division site once the Z ring is assembled. Together, these results suggest that HVO_0399 plays an important role in *H. volcanii* cell division, so we renamed it <u>C</u>ell <u>D</u>ivision <u>P</u>rotein <u>C</u> (CdpC).

**Fig. 2.**
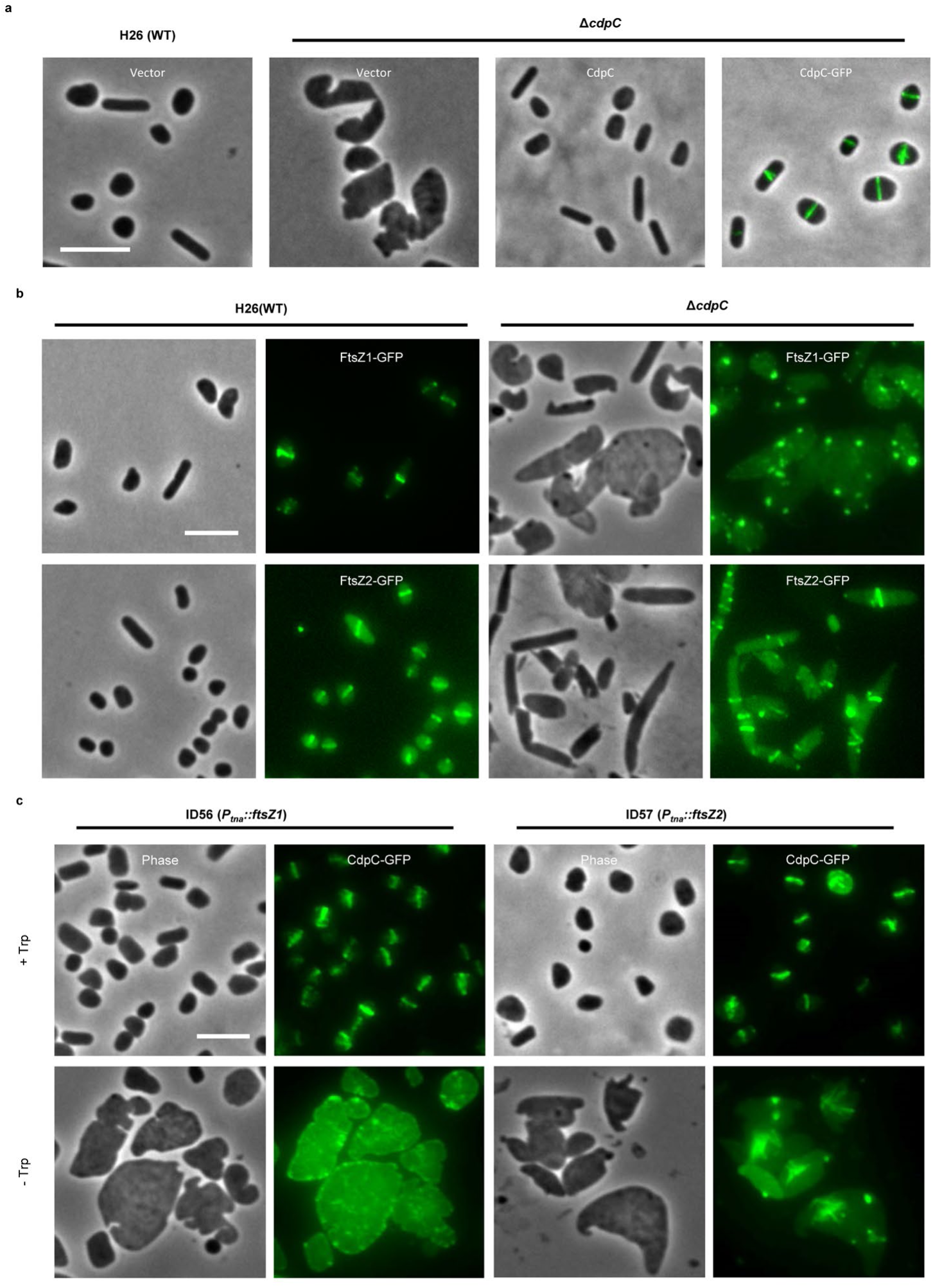
CdpC is important for FtsZ1 ring formation and cell division in *H. volcanii*. a. Representative images *ΔcdpC* cells complemented with CdpC or its GFP fusion expressed from a plasmid. Cells were cultured in Hv-Cab medium at 45 °C, expression of CdpC and the CdpC-GFP fusion were driven by their native promoter. b. Representative images of the localization FtsZ1-GFP or FtsZ2-GFP in wild type and *ΔcdpC* cells. Cells were cultured in Hv-Cab medium at 45 °C, FtsZ1-GFP and FtsZ2-GFP were expressed from their native promoters from plasmids. c. Representative images of the localization of CdpC-GFP in FtsZ1 or FtsZ2 depleted cells. Cells were cultured in Hv-Cab medium at 45 °C, expression of CdpC was driven by its native promoter. Depletion of FtsZ1 or FtsZ2 was achieved by removal of tryptophan (Trp) from the cultures. Scale bars in a-c, 5 μm.

To determine why the absence of CdpC results in severe division and shape defects, we examined the localization of key cell division proteins in *ΔcdpC* cells. Strikingly, FtsZ1-GFP was diffusely distributed and formed bright foci in the cytoplasm in *ΔcdpC* cells, in sharp contrast to the ring-like localization in wild type cells (Fig. 2b). However, GFP fusions to FtsZ2 and the other division proteins (SepF, CdpA, CdpB1/2/3) retained their ability to form filaments and spirals (Fig. 2b, Extended Data Fig. 2b), even though they were often mis-localized in these abnormal cells. that the absence of CdpC prevents FtsZ1 from assembling into the Z ring, subsequently leading to the severe division and shape defects. Thus, CdpC is essential for FtsZ1 ring formation but is not required for the assembly of FtsZ2 and its associated proteins.

Next, we determined how CdpC is recruited to the midcell by examination of its localization in strains lacking individual key cell division proteins, including FtsZ1, FtsZ2, SepF, CdpA and CdpB1. We expressed CdpC-GFP in deletion mutants or depletion strains which had the gene to be tested under the control of the *P_tna_* promoter. As expected, CdpC-GFP localized to the midcell in wild type cells and in all the depletion strains in the presence of tryptophan (Fig. 2c, Extended Data Fig. 3). However, in the absence of tryptophan, CdpC-GFP displayed a diffusive and membrane-associated distribution in FtsZ1 depleted cells (Fig. 2c), indicating that it depends on FtsZ1 for proper localization. Although CdpC-GFP could not form typical ring-like structures in the irregular and giant cells after depletion of FtsZ2, SepF, CdpB1, or deletion of *cdpA*, it was present in filamentous structures in these cells (Fig. 2c, Extended Data Fig. 3), presumably by association with FtsZ1. Together, these results suggest that CdpC depends on FtsZ1, but not other division proteins, to localize to the midcell.

### CdpC interacts directly with FtsZ1

The interdependent localization pattern between CdpC and FtsZ1 prompted us to determine whether they interact with each other directly. To do this, we first used the Split-FP (fluorescent protein) assay, which is based on the reconstitution of the superfolder GFP upon proximity of two target proteins^26^. Strong fluorescent signals and fluorescent rings were observed in cells co-expressing tagged versions of CdpC with FtsZ1, but not in cells expressing only the tags or when only one of the proteins was tagged (Fig. 3a,b). We also detected an interaction signal between CdpC and FtsZ2 and observed midcell localization of the reconstituted sfGFP (Extended Data Fig. 4a,b). However, the higher fluorescence signal observed with FtsZ1 suggests a stronger or more specific interaction between CdpC and FtsZ1.

**Fig. 3.**
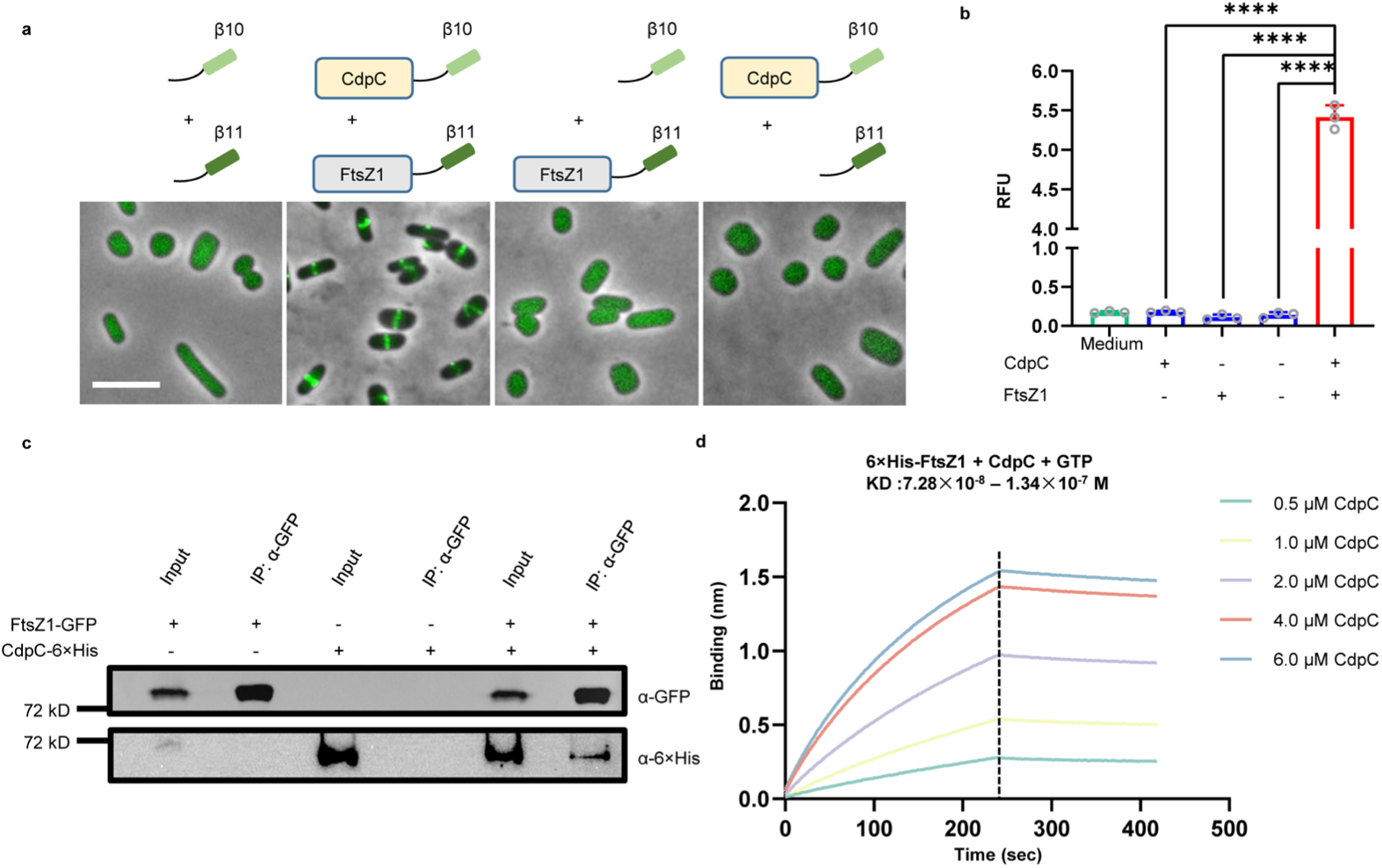
CdpC interacts directly with FtsZ1 *in vivo* and *in vitro*. a. Representative images of the fluorescence of Split-FP assay to assess the interaction between CdpC and FtsZ1 *in vivo*. Scale bar, 5 μm. b. Quantitation of the interaction signal between CdpC and FtsZ1 in the Split-FP assay in H26 (WT). RFU, relative fluorescence unit. Data are presented as mean values ± s.d. Significance in each group was tested by two-sided t-test. ****P <0.0001, n=3. c, Co-IP experiment showing that CdpC interacts with FtsZ1 *in vivo*. Cultures of *H. volcanii* expressing the indicated proteins were lysed by sonication; supernatants were incubated with anti-GFP antibodies coated magnetic beads. Immunocomplexes were eluted with boiling SDS-PAGE loading buffer and then analyzed by immunoblot. d, BLI assay showing that CdpC interacts with FtsZ1 *in vitro*. Details about the experimental set up and procedures are described in Methods. The concentration of the 6×his-FtsZ1 protein remains constant (0.5 μM), the concentration of GTP is 1 mM, while the concentration of CdpC varies.

To confirm the interaction between CdpC and FtsZ1, we performed Co-IP experiments in cells co-expressing CdpC-6×His and FtsZ1-GFP. As expected, the two proteins co-precipitated (Fig. 3c, Extended Data Fig. 4c). In contrast, CdpC-6×His was not detected in the immunoprecipitates from cells co-expressing CdpC-6×His and FtsZ2-GFP (Extended Data Fig. 4d,e). These results indicate that CdpC interacts directly with FtsZ1, but not with FtsZ2, *in vivo*. To further validate the interaction, we purified CdpC using the SUMO purification system and 6×His-FtsZ1 and tested their interaction by bio-layer interferometry (BLI) assays. As shown in Fig. 3d, CdpC bound to 6×His-FtsZ1 robustly with a dissociation constant (KD) ranging from 7.28 × 10⁻⁸ to 1.34 × 10⁻^7^ M. Altogether, these results demonstrate that CdpC binds directly to FtsZ1 with high affinity.

### CdpC possesses three distinct functional domains

To further characterize the function of CdpC, we used AlphaFold 3 (AF3)^27^ to generate a structural model, which revealed three primary domains: an N-terminal domain (NTD, residues 1–60), a C-terminal domain (CTD, residues 304–346), connected by a long intrinsically disordered linker (residues 61–303) (Fig. 4a, and Extended Data Fig. 5a). Interestingly, both the NTD and CTD of CdpC were predicted with high confidence to exhibit a zinc ribbon architecture, each contained a zinc-finger motif (Extended Data Fig. 5b,c,e,f). Note that since CdpC is distant (<10% sequence homology) from any proteins of known function, the disordered linker may also contain a more structured domain failed to be predicted by AF3. To determine the function of each domain of CdpC, we checked the localization of GFP fusions in wild type cells. Strikingly, the CTD (CdpC_CTD_-GFP) localized to midcell as well as full length CdpC-GFP, whereas the NTD (CdpC_NTD_-GFP) and the linker (CdpC_Linker_-GFP) appeared to associate with the membrane and be diffusely distributed in the cytoplasm, respectively (Fig. 4b). These results suggest that CdpC_NTD_ binds to the membrane, while CdpC_CTD_ likely interacts with FtsZ1. In support of this, Split-FP assays showed strong fluorescent signals and midcell rings of sfGFP in cells co-expressing tagged versions of FtsZ1 and CdpC_CTD_ (Extended Data Fig. 6a-f). It is notable that cells co-expressing tagged versions of FtsZ1 and CdpC_NTD_ exhibited weak fluorescence and rare midcell rings, indicating that CdpC_NTD_ may also interact with FtsZ1 or be in close proximity to it by association with the divisome. In contrast, cells co-expressing tagged versions of FtsZ1 and CdpC_Linker_ yielded no detectable fluorescence signal and ring (Extended Data Fig. 6a-d). Taken together, these results indicate that CdpC_NTD_ associates with the membrane and CdpC_CTD_ interacts strongly with FtsZ1, whereas CdpC_Linker_ connects these two functional modules.

**Fig. 4.**
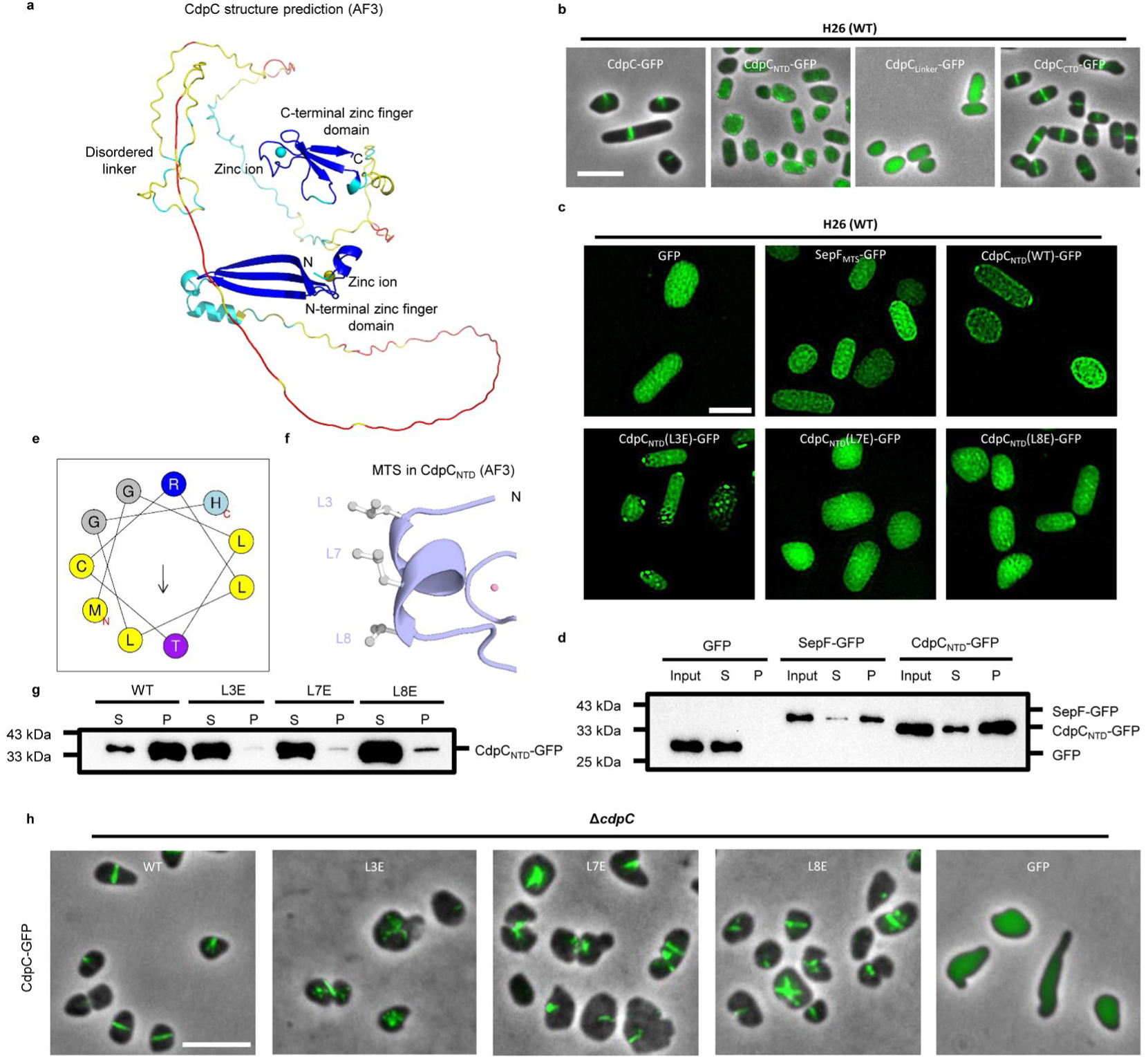
Structural model of CdpC and its NTD mediates membrane binding. a. Alphafold 3 structural model of CdpC. Both its NTD and CTD contain a zinc finger. Model confidence (blue - pLDDT > 90%, cyan - 90% > pLDDT > 70%, yellow – 70% > pLDDT > 50%, red - pLDDT <50%), ipTM = 0.46, pTM=0.23. Zinc ions are indicated. b. Representative images of the localization of CdpC_NTD_-GFP, CdpC_Linker_-GFP, or CdpC_CTD_-GFP in H26 (WT) cells. Scale bar, 5 μm. c. Representative images of the localization of CdpC_NTD_-GFP and its mutant forms by 2D-SIM. GFP and GFP fusion proteins were expressed from plasmids under the control of the *P_tna_* promoter, induced overnight with 1 mM tryptophan (Trp) at 30 °C. Scale bar, 2 μm. d. Membrane fractionation assay showing that the NTD of CdpC binds to the cell membrane *in vivo*. Cultures of *H. volcanii* expressing the indicated proteins were lysed by sonication; the membrane fraction was isolated by ultracentrifugation of the supernatant of lysed cells; the pellet was resuspended to the original volume and analyzed by immunoblot. S, supernatant; P, pellet. e. HeliQuest analysis of the N-terminal helix of CdpC. Color code: yellow (non-polar), purple (polar), blue (charged), gray (Gly). Arrow points to the hydrophobic face. f. The membrane targeting sequence (MTS) in CdpC_NTD_ as it appears in the Alphafold3 structure model. Atom colors: gray, carbon; pink, zinc ion. g. Membrane fractionation assay showing the effect of the mutations in the amphipathic helix of CdpC on membrane association *in vivo*. The procedure for the test was similar to d. S, supernatant; P, pellet. h. Representative images of the localization of CdpC-GFP and its mutants in *ΔcdpC* cells. Cells were cultured in Hv-Cab medium at 45 °C, GFP and GFP fusion proteins were expressed from plasmids under the control of the P_tna_ promoter, induced overnight with 0.2 mM tryptophan (Trp). Scale bar, 5 μm.

### CdpC_NTD_ associates with the cell membrane with an amphipathic helix

To confirm that CdpC_NTD_ associates with the membrane, we first used structured-illumination microscopy (SIM)^28^ to observe its subcellular localization in wild type cells. In contrast to the diffusive cytoplasmic distribution of GFP, CdpC_NTD_-GFP exhibited a clear membrane association, similar to the membrane-targeting sequence of SepF (SepF-MTS) (Fig. 4c). Ultracentrifugation-based membrane fractionation showed that CdpC_NTD_-GFP, similar to SepF-GFP, was predominantly enriched in the pellet (membrane fraction), while GFP was primarily in the supernatant (cytosol) (Fig. 4d). These results strongly indicate that CdpC_NTD_ is able to bind to the membrane.

To determine how CdpC_NTD_ binds to the membrane, we used HeliQuest ^29^ to analyze its primary sequence and found that it contains an amphipathic helix at the extreme N-terminus (Fig. 4e), which can target a cytoplasmic protein to the membrane. In line with this, there is an α-helix in the structural model of CdpC_NTD_ with three hydrophobic leucine residues clustered on one face (Fig. 4f). To test whether these leucine residues are critical for membrane anchoring, we mutated each to glutamate (L3E, L7E and L8E) and examined their impact on the localization of CdpC_NTD_-GFP. As expected, all three mutations abolished the membrane localization of CdpC_NTD_-GFP: the L3E mutant was not at the membrane and formed some bright foci in the cytoplasm, while L7E and L8E mutants displayed a diffusive cytoplasmic distribution similar to that of GFP (Fig. 4c). Membrane fractionation assays showed that all three mutants of CdpC_NTD_-GFP were predominantly enriched in the supernatant (cytosol) (Fig. 4g). Western blotting showed that none of the mutations affected the stability of CdpC_NTD_ or full-length CdpC (Extended Data Fig. 7a,b).

To test if membrane binding is important for CdpC function, these mutations were introduced into full-length CdpC-GFP. Complementation tests and localization analysis revealed that all three mutations abolished the ability of the fusion protein to form normal rings and restore normal cell division and shape in Δ*cdpC* cells (Fig. 4h). However, the fluorescence was present in filamentous structures suggesting these mutants associated with FtsZ1 filaments. In summary, these results demonstrate that the amphipathic helix embedded in CdpC_NTD_ mediates the membrane association of CdpC and is critical for its function.

### CdpC_CTD_ interacts directly with FtsZ1 and promotes its assembly *in vivo*

To confirm the interaction between CdpC_CTD_ and FtsZ1, we performed Co-IP assays with tagged versions of CdpC_CTD_ and FtsZ1. As expected, CdpC_CTD_-GFP co-precipitated with FtsZ1-6×His from cells in which they were co-expressed (Fig. 5a,b). To further validate the interaction, we synthesized 6×His-CdpC_CTD_ and purified FtsZ1 and tested their interaction by BLI assays. As shown in Fig. 5c, FtsZ1 bound to 6×His-CdpC_CTD_ robustly, with a KD value ranging from 7.42 × 10⁻⁸ M to 1.20 × 10⁻^7^ M, similar to the KD value between full length CdpC and FtsZ1. These results demonstrate that CdpC_CTD_ interacts directly with FtsZ1.

**Fig. 5.**
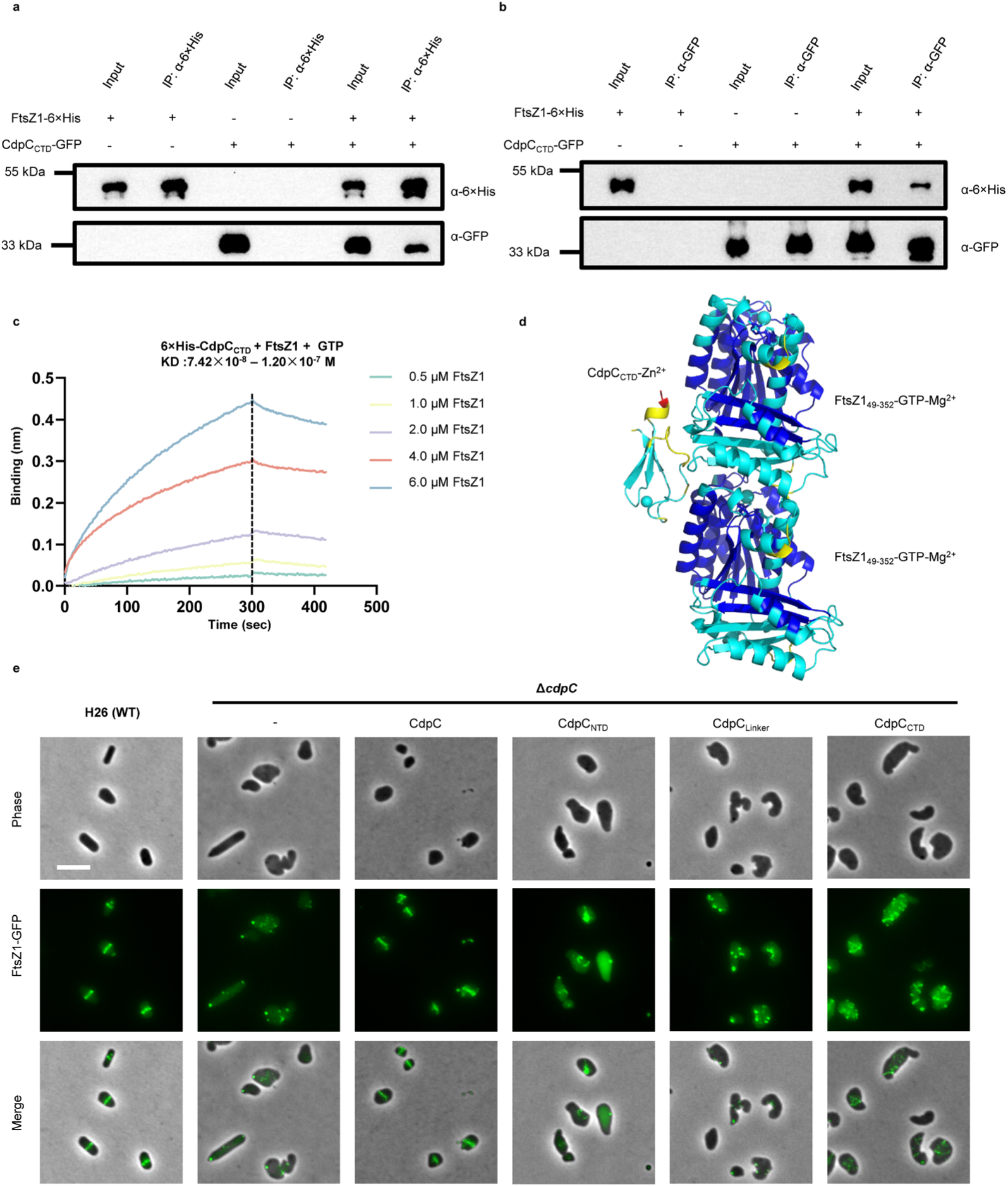
The CTD of CdpC interacts directly with FtsZ1. a-b. Co-IP experiments showing that CdpC_CTD_ interacts with FtsZ1 *in vivo*. Cultures of *H. volcanii* expressing the indicated proteins were lysed by sonication and supernatants from the lysates were incubated with anti-GFP (a) or anti-6×his (b) antibodies coated magnetic beads. Immunocomplexes were eluted from the beads with boiling SDS–PAGE loading buffer and then analyzed by immunoblot. c. BLI assay showing that CdpC_CTD_ interacts with FtsZ1 *in vitro*. Details about the experimental set up and procedures are described in Methods. The concentration of the 6×his-CdpC_CTD_ remains constant (0.5 μM), the concentration of GTP is 1 mM, while the concentration FtsZ1 of varies. d. A structural model of the CdpC_CTD_-FtsZ1 complex at a stoichiometry of 1:2 of CdpC_CTD_ to FtsZ1_49-352_ with zinc ion, magnesium ion and GTP. Model confidence (blue - pLDDT > 90%, cyan - 90% > pLDDT > 70%, yellow - 70% > pLDDT > 50%, red - pLDDT <50%), ipTM = 0.45, pTM= 0.54. e, Representative images of the localization of FtsZ1-GFP in *ΔcdpC* cells expressing one of the three domains of CdpC. Cells were cultured in Hv-Cab medium at 45 °C, FtsZ1-GFP and CdpC (or its various domains) were expressed from the same plasmid, with FtsZ1-GFP driven by its native promoter and CdpC (or its individual domain) under the control of the *P_tna_* promoter. Cultures were induced with 0.2 mM tryptophan (Trp) overnight before imaging. Scale bar, 5 μm.

To explore how CdpC_CTD_ interacts with FtsZ1, we generated structural models for their complex (in the absence of the N-terminal tail, linker and C-terminal tail of FtsZ1) with different stoichiometries using AF3 ^27^. With a 1:2 stoichiometry of CdpC_CTD_/ FtsZ1_49-352_, we obtained a model with modest confidence which suggests that CdpC_CTD_ binds to the exposed surface of the interface between two FtsZ1_49-352_ molecules (Fig. 5d, and Extended Data Fig. 5d). This implied that CdpC promotes FtsZ1 polymerization and was consistent with the observation that FtsZ1 failed to assemble into filaments and rings in the absence of CdpC (Fig. 2b). To test this, we examined FtsZ1-GFP localization in *ΔcdpC* cells complemented with different domains of CdpC. Neither CdpC_NTD_ nor CdpC_Linker_ could restore FtsZ1-GFP ring formation in *ΔcdpC* cells as it was diffusely distributed and formed bright foci in the cytoplasm (Fig. 5e). However, in *ΔcdpC* cells expressing CdpC_CTD_, numerous disorganized FtsZ1 filaments or spirals were observed, although midcell FtsZ1-GFP ring was not restored (Fig. 5e). These results agree with the structural model suggesting that CdpC_CTD_ facilitates FtsZ1 polymerization by binding to the exposed surface of FtsZ1 subunit interfaces.

### The zinc fingers of CdpC are essential for its function

CdpC is predicted to be a zinc finger protein with both its NTD and CTD containing a zinc finger followed by three β-strands, forming zinc ribbons. Zinc finger proteins have been extensively studied in eukaryotes and bacteria, however, their function in archaea have not been clearly defined^30^. Zinc ions can be coordinated by zinc fingers containing four cysteine (C4), three cysteine and one histidine (C3H), or two cysteine and two histidine (C2H2)^30^. The zinc-finger in CdpC_NTD_ is a C3H type (C5, H10, C38 and C41), while the one in CdpC_CTD_ belongs to type C4 (Fig. 6a,b). Substitution of any one of the four putative zinc-coordinating residues in CdpC_NTD_ with alanine (C5A, H10A, C38A, C41A) did not affect the protein level of full-length CdpC-GFP, however, they completely destabilized CdpC_NTD_-GFP for unknown reasons (Extended Data Fig. 7c,d). Thus, we assessed the impact of the mutations on CdpC function using full-length CdpC-GFP. As shown in Fig. 6c, these mutations severely impaired the ability of CdpC-GFP to complement *ΔcdpC* cells. Moreover, the mutants were present in disorganized filaments, similar to CdpC_CTD_-GFP, suggesting that membrane binding of CdpC was disrupted by the mutations. However, in wild-type cells the CdpC-GFP mutants still localized to the mid-cell, presumably because endogenous CdpC supports FtsZ1 ring assembly and the mutants’ CTD is sufficient to associate with FtsZ1 (Extended Data Fig. 8a). Notably, one of the zinc-coordinating cysteine (C5) is located within the N-terminal amphipathic helix, implying that disruption of the zinc finger may destabilize the amphipathic helix, consequently compromising membrane association of CdpC.

**Fig. 6.**
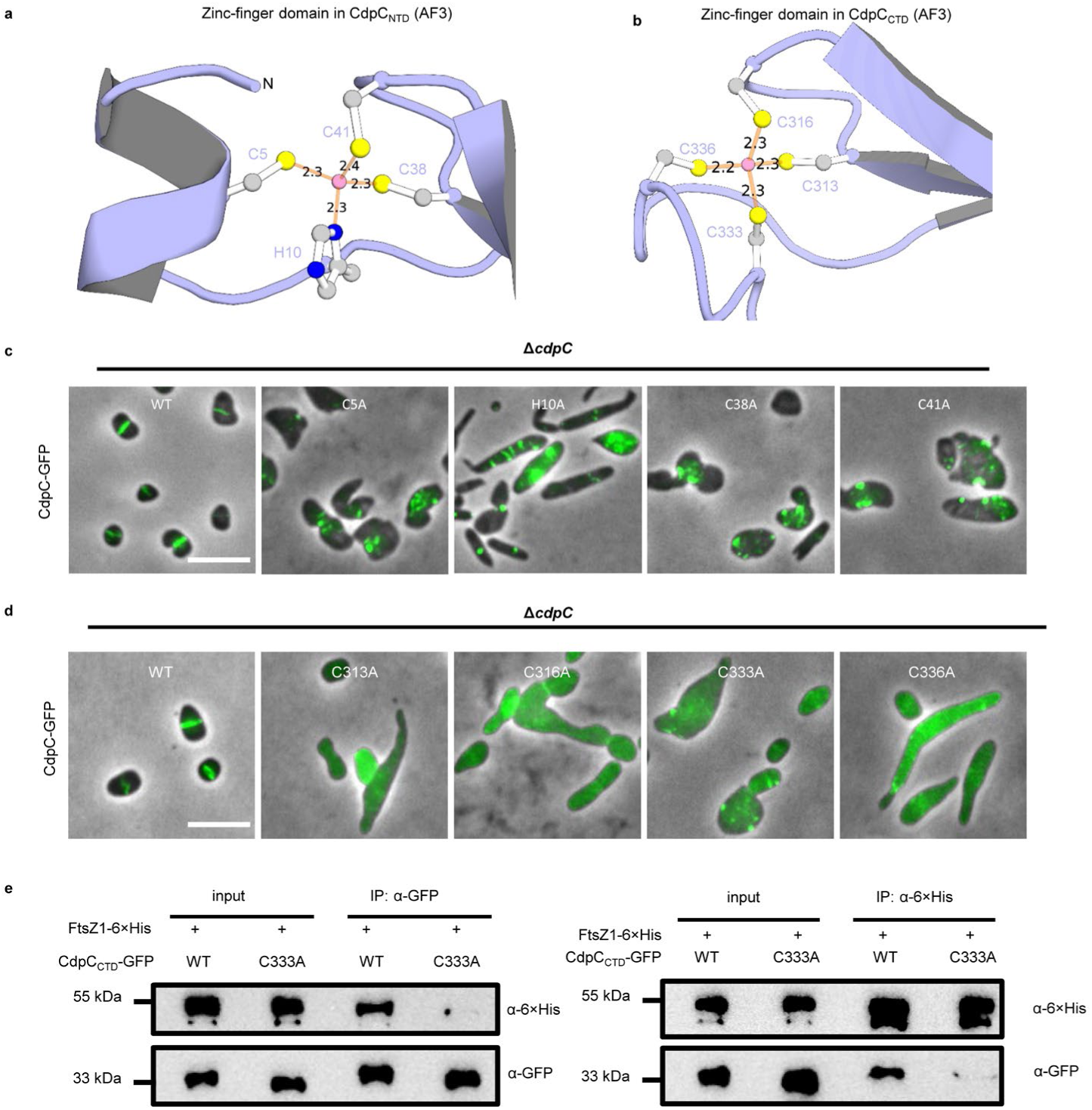
Zinc finger motifs in the NTD and CTD of CdpC are important for its functions. a-b. Zinc finger motifs in CdpC_NTD_ (a) and CdpC_CTD_ (b) derived from the Alphafold 3 structural model of CdpC. Residues coordinating the zinc ion are shown. Atom colors: gray, carbon; yellow, sulfur; blue, nitrogen; pink, zinc ion. Coordination bonds are shown as orange lines, with distances (Å) labeled. c-d, Representative images of the localization of CdpC-GFP mutants with mutations in the zinc finger in CdpC_NTD_ or in CdpC_CTD_ in *ΔcdpC* cells. Cells were cultured in Hv-Cab medium at 45 °C, GFP fusion proteins were expressed from plasmids under the control of the *P_tna_* promoter, induced overnight with 0.2 mM tryptophan (Trp). Scale bars, 5 μm. e, Co-IP experiment showing that a mutation in the zinc-binding site disrupts the interaction between CdpC_CTD_ and FtsZ1. Cultures of *H. volcanii* expressing the indicated proteins were lysed by sonication; supernatants were incubated with anti-GFP or anti-6×his antibodies coated magnetic beads. Immunocomplexes were eluted with boiling SDS-PAGE loading buffer and then analyzed by immunoblot.

We also substituted the putative zinc coordinating residues in CdpC_CTD_ and full length CdpC. None of the substitutions significantly affected protein stability (Extended Data Fig. 8e,f), however, all four mutations disrupt function. In either wild type H26 or *ΔcdpC* cells, CdpC_CTD_-GFP mutants failed to localize at midcell and showed diffusive cytoplasmic distribution (Extended Data Fig. 8c,d). Similarly, full-length CdpC-GFP carrying zinc-coordinating mutations in the CTD lost midcell localization and exhibited membrane-associated localization, similar to CdpC_NTD_ (Fig. 6d, Extended Data Fig. 8b). These results indicate that disruption of the zinc finger in CdpC_CTD_ abrogates its ability to interact with FtsZ1, thereby preventing its midcell localization and proper function. To validate this, we examined the interaction between FtsZ1 and CdpC_CTD_ zinc-coordinating mutants via the Split-FP assay. When tagged versions of wild type CdpC_CTD_ and FtsZ1 were co-expressed, strong fluorescence signals and clear midcell rings were observed (Extended Data Fig. 9a, b). In contrast, co-expression of any of the four CdpC_CTD_ mutants with FtsZ1 yielded little fluorescence signal (Extended Data Fig. 9a, b), indicating a loss of interaction. Moreover, Co-IP in cells co-expressing FtsZ1-6×His and one representative CdpC_CTD_-GFP mutant, C333A, contained only trace amounts of the mutant in the immunoprecipates compared to wild type (Fig. 6e). Altogether, these results indicate that zinc-binding is critical for CdpC_CTD_ to interact with FtsZ1.

### CdpC-like proteins evolved fast but maintained conserved zinc ribbon domains

Having uncovered the role of CdpC in *H. volcanii* division, we explored its distribution across different lineages of archaea. Initial Blast search using *H. volcanii* CdpC (*Hv*CdpC) only fetched sequences almost exclusively from the *Haloferacaceae* family, among which shared amino acid identity could drop below 30%. Given the essential role of *Hv*CdpC in cell division, we expected that it would have homologs in a broader taxonomic range. We thus carried out an iterative search process based on Hidden Markov Models (HMMs)^31^ (See Methods). This approach led us to find that CdpC-related proteins with similar NTD-linker-CTD domain architecture broadly encoded across the *Halobacteria* class (in 617 out of 631 genomes, Fig. 7a). Phylogenetics indicated that CdpC is vertically transmitted but evolved fast, showing only around 20% sequence similarity between different families within *Halobacteria*. However, Multiple Sequence Alignment (MSA) indicates that the above identified N-terminal amphipathic helix with hydrophobic Leucine residues, as well as the zinc finger motifs in the NTD and CTD are most conserved (Fig. 7b). By contrast, the linker region is highly variable, contributing to the steep drop in sequence similarity across species evolutionary distances. These features were also well recovered via AF3 predictions (Extended Data Fig. 10a). However, we do observe an enrichment in amino acid residue conservation at positions 207-253 within the linker of *Hv*CdpC (Fig. 7b). It is possible that this represents a structured motif failed to be predicted by AF3. Overall, Halobacterial CdpC exhibited a strong conservation in structure as well as key binding residues, albeit highly variable sequences, implicating a conserved mechanism as an FtsZ membrane anchor across *Halobacteria*.

**Fig. 7.**
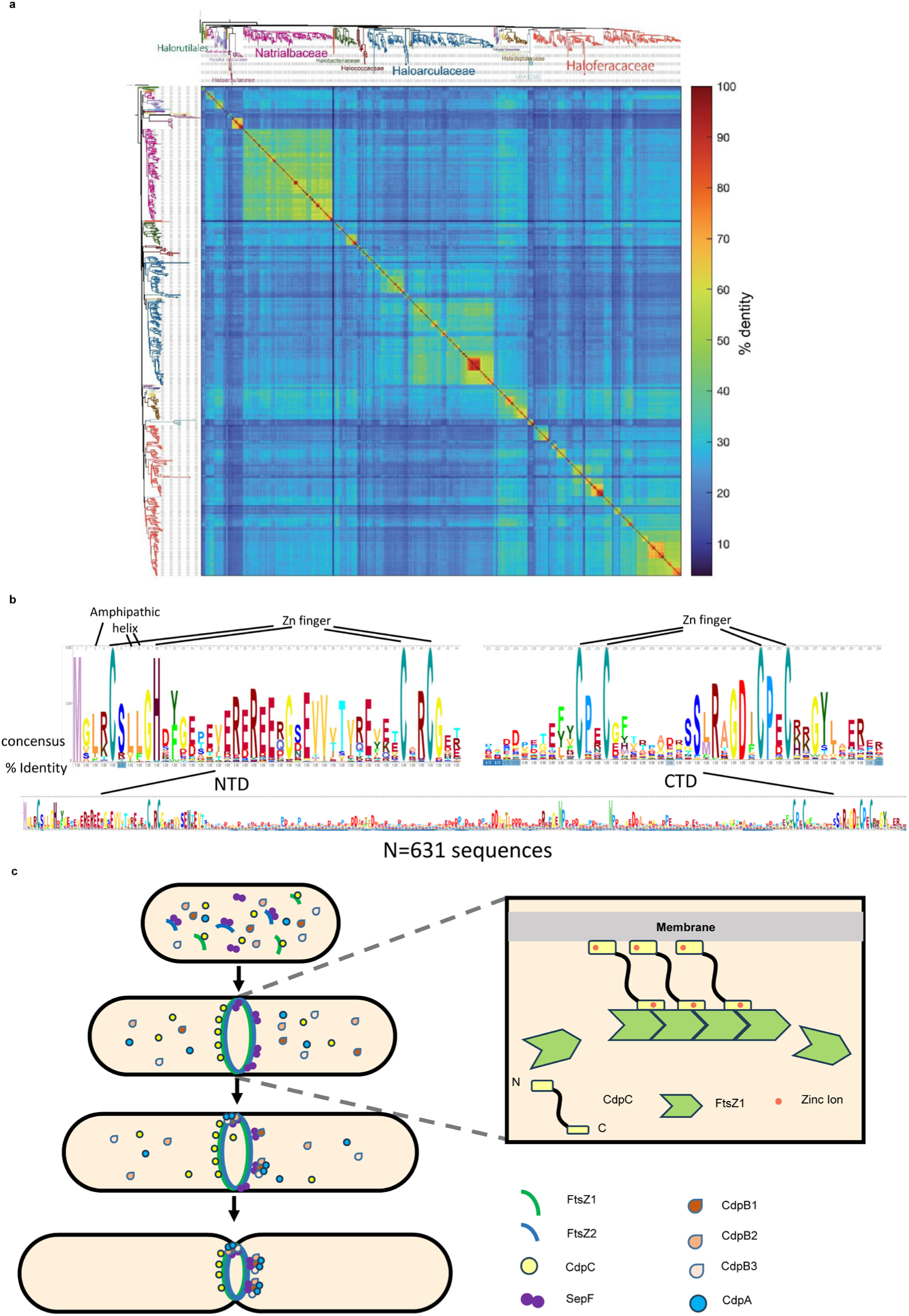
Conservation of CdpC and a working model for its function in *H. volcanii* cell division. a. Pairwise amino acid identity matrix of 631 CdpC homologs across 617 Halobacterial species. The corresponding phylogenetic tree is indicated on top and left, where different families of *Halobacteriales* and the less-sampled order *Halorutilales* are indicated in different colors on top. Color bar indicates the % amino acid identity in the matrix. b. Consensus sequences of the 631 CdpC homologs across 617 Halobacterial species show the conserved Leucine residues in the amphipathic helix (top left), and the Zinc-interacting residues in the two zinc finger motifs. c. A working model of CdpC function in *H. volcanii*. CdpC acts as a membrane anchor for FtsZ1 and promotes FtsZ1 assembly into the Z1-ring at the initiation of cytokinesis. The NTD and CTD of CdpC bind to the membrane and FtsZ1, respectively. Zinc binding is important for CdpC function. Subsequently, FtsZ2, SepF, CdpA and CdpB1-3 join the Z1-ring and recruit downstream cell division proteins to complete divisome assembly, leading to cytokinesis.

Considering the fast-evolving nature of the CdpC protein family, we next explored whether there are proteins with remote homology that exhibit similar structural features by extending the iterative HMM process towards more divergence sequences. To this end, we found sequences in all classes of Thermoplasmatota, in most DPANN phyla (including Nanobdellota, Micrarchaeota, EX4484-52, Altiarchaeota, and Aenigmatarchaeota), as well as in Lokiarchaeia within the Asgard archaea clade. AF3 structural prediction indicates the existence of multiple zinc ribbon domains across these proteins, analogous to *Hv*CdpC with double zinc ribbons (Extended Data Fig. 10b). The unstructured linker region is, however, not apparent in these proteins. Notably, no homolog was found within the Thermoproteota except for one singleton, coinciding with the general lack of FtsZ-based cell division in this phylum. Overall, multi-zinc-ribbon proteins distantly related to *Hv*CdpC are widespread in archaea, and may contribute to cell division.

## Discussion

Compared to bacteria and eukaryotes, the mechanisms underlying archaeal division remain poorly understood. In this study, we discovered ten candidate division proteins in *H. volcanii* by *in vivo* CLIP-MS. Among these, CdpC functions as a membrane anchor for FtsZ1 and is important for cell division (Fig. 7c). Specifically, its NTD associates with the membrane via an amphipathic helix, while its CTD interacts directly with FtsZ1 and promotes its assembly into the Z ring. Notably, both the NTD and CTD of CdpC are zinc ribbon structures containing a zinc finger motif that is essential for its function. Based on our findings, we speculate that CdpC promotes FtsZ1 polymerization and tethers FtsZ1 filaments to the membrane to form the FtsZ1 ring to initiate divisome assembly in *H. volcanii*. Subsequently, FtsZ2 and SepF join the ring, followed by CdpA and CdpB1/2/3. These proteins assemble into the early divisome which likely recruits additional division proteins to form the complete divisome, ultimately leading to cell constriction (Fig. 7c).

Formation of a Z ring is the initial step of cytokinesis in prokaryotes relying on FtsZ for division^32^. Bacteria employ diverse proteins to anchor FtsZ filaments to the membrane^33^. Some of these membrane anchors, such as FtsA, which is an actin-like protein, and SepF, are widely conserved and can assemble into polymers bound to the membrane through an amphipathic helix^33^. Others, such as EzrA and ZipA, are less-conserved and contain transmembrane domains^33^. SepF is the only known membrane anchor for FtsZ in archaea before this study, but it specifically anchors FtsZ2 to the membrane in archaea containing two FtsZs. Given that FtsZ1 is believed to play a predominant role in forming the Z ring^11,17^, it has been a long-standing puzzle how it is anchored to the membrane. Here we discover that CdpC anchors FtsZ1 filaments to the membrane via an NTD with an amphipathic helix and a CTD that binds to FtsZ1 directly in *H. volcanii*, resolving a critical question in divisome assembly. However, unlike most of the FtsZ membrane anchors which bind to the conserved C-terminal tail of FtsZ, CdpC seems to bind to the globular domain of FtsZ1. Archaea and bacteria thus appeared to have convergently evolved amphipathic helix-based Z-ring membrane anchors, while using drastically different structural basis to interact with the Z-ring.

Although our study reveals an important role of CdpC in *H. volcanii* cell division, its detailed working mechanism warrants further investigation. First, the localization pattern and Split-FP data hint at a weak or indirect interaction between its NTD and FtsZ1. It is possible that the NTD directly interacts with FtsZ1 or interacts with other divisome proteins beyond mediating the membrane association of CdpC. Second, although our data and the AF3 structural model suggest that CdpC_CTD_ binds to FtsZ1 directly and facilitates its polymerization, how exactly it works remains to be elucidated. Mutational analysis and structural information are necessary to validate the interaction mode between CdpC_CTD_ and FtsZ1. Third, whether the extraordinarily long 243-amino-acid linker domain, which appeared unstructured in AF3, possesses functions beyond providing a mere physical connection, such as recruiting downstream division proteins, merits exploration. Lastly, in addition to serving as the FtsZ1 membrane anchor, whether CdpC plays a role in recruitment of other divisome proteins awaits investigation.

It is notable that CdpC contains two zinc ribbon domains and disrupting either of the zinc finger motifs in these domains perturbs cell division. Proteins with zinc finger motifs perform a variety of functions in eukaryotic and bacterial cells, including DNA binding as transcription factors, and mediating interactions with RNA, proteins and lipids^34^. Recent studies found that proteins with zinc finger motifs are also abundant in archaea and *H. volcanii* harbors 49 putative zinc finger proteins with two CPXCG motifs^30,34^. These proteins have been implicated in transcriptional regulation, stress adaptation, motility, and biofilm formation^30^. However, here we find that CdpC, harboring a dual zinc ribbon architecture, plays a critical role in *H. volcanii* cell division instead of the above suggested functions. Indeed, distant homologs of CdpC outside Halobacteria are multiple zinc ribbon-containing proteins with unknown function, and are absent in Thermoproteota, which do not use FtsZ-based cell division system. It would be interesting in the future to examine the possible interactions between these proteins and FtsZ1 in their respective organisms and their roles in archaeal cell division.

In addition to CdpC, this study reports an additional nine proteins that display clear midcell localization. Except for HVO_0792, none of these proteins appears to strongly affect division or morphology under standard laboratory conditions. However, this does not necessarily mean that they are not involved in cell division. Absence of phenotype in single gene deletions may be caused by compensatory effects between different components of the divisome, as observed in bacteria. Alternatively, it is also plausible that many of them are accessory divisome proteins, playing important roles in cell division under specific stress conditions, or in other archaeal groups, similar to accessary proteins of the bacterial divisome. Strikingly, about half of these proteins are putative metabolic enzymes, indicating a coupling of cell division and metabolism in haloarchaea. HVO_0792 is a homolog of the shikimate pathway enzyme DHQS, while HVO_2958, HVO_2959, HVO_2960 and HVO_2961 are subunits of the OADHC complex. These proteins are either moonlighting enzymes involved in both metabolism and cell division or taking advantage of the divisome for proper localization to enhance their efficiency. The precise connections and functional contributions of these proteins to cell division and metabolism present promising avenues for future research.

In summary, this work significantly increases our understanding of the composition of the divisome and uncovers the mechanism of Z ring formation in haloarchaea. Also, this work suggests that the abundant zinc finger proteins in archaea—akin to eukaryotes—may play critical roles in diverse cellular processes. Additionally, there may be a delicate coupling mechanism between cell division and metabolism in archaea.

## Methods

### Strains, plasmids, and growth conditions

All strains and plasmids used in this study are listed in Supplementary Table 4 and Table 5, respectively. *Escherichia coli* strains JS238, JM110 and BL21(DE3) were used for plasmid propagation, de-methylation and protein expression, respectively. Cells were grown in Luria-Bertani (LB) broth (10 g/L tryptone, 5 g/L yeast extract, 5 g/L NaCl, and 0.05 mg/mL of thymine) or on LB agar plates (1.5% agar) at indicated temperatures. When required, ampicillin was added to a final concentration of 100 μg/mL to maintain plasmids. *H. volcanii* strains were routinely cultured at 45 °C in Hv-YPCTE or Hv-Cab medium as described previously^19,35^. When required, 5-fluoroorotic acid (FOA, 50 μg/mL) was added for counter selection. L-Tryptophan (Trp) was added at indicated concentrations to induce protein expression from the P*_tna_* promoter. Growth was monitored by optical density at 600 nm (OD_600_); cultures were grown in exponential phase (OD_600_< 0.8) before sampling unless stated otherwise.

### Reagents and chemicals

All reagents and chemicals are listed in Supplementary Table 7.

### Genomic modifications

Gene deletion and promoter-replacement strains were constructed using a two-step homologous recombination (pop-in/pop-out) strategy^24^. To construct the deletion strain of *cdpC* in *H. volcanii*, the de-methylated non-replicating plasmid carrying upstream and downstream flanking regions of the start codon of *cdpC* were introduced into *H. volcanii* H26 by PEG-mediated spheroplast transformation. Single-crossover integrants were selected on uracil-free Hv-Cab agar, followed by counterselection on FOA-containing Hv-YPCTE plates to excise the plasmid. The desired chromosomal alterations were verified by PCR and Sanger sequencing. The same approach was applied to construct deletion strains for *hvo_1130*, *hvo_1708*, *hvo_1512*, *hvo_1301*, *hvo_2958*, *hvo_2959*, *hvo_2960*, *hvo_2961*, and *cdpA* in *H. volcanii* H26. A depletion strain of *hvo_0792* was constructed by the same approach with the non-replicating plasmid carrying the upstream flanking region of *hvo_0792* and L11e transcription terminator followed by the *P_tna_::hvo_0792* cassette in *H. volcanii* H26.

### Plasmid construction

All plasmids used in this study are listed in Supplementary Table 5 and were constructed using the primers in Supplementary Table 6. Plasmids used for genomic modification, controlled gene expression and split-GFP assays in *H. volcanii* were constructed with the parental plasmids pTA131^24^, pTA962 and pTA1228^36^, respectively. Plasmids used for complementation assays in the *ΔcdpC* strain expressed variants of CdpC under the control of its native promoter. GFP or mCherry fusions of FtsZ1, FtsZ2, SepF and CdpB1 were expressed under their respective native promoters. For all other genes, GFP fusions were driven by the *P_tna_* promoter, except when expressed in depletion strains, in which case their native promoters were used instead. To construct GFP or mCherry fusions, the coding sequences of target genes were amplified and inserted in-frame with the reporter gene. The *P_tna_* promoter of the above plasmids was replaced by the native promoters of the respective genes for use in depletion strains. To construct dual expression plasmids of the various genes in *H. volcanii*, a fragment containing one of the genes was ligated into the NotI-cut site of the above plasmids (which already contained the expression cassette for the other gene). For split-GFP assays, the sfGFP10 and sfGFP11 fragments were fused to the proteins of interest via flexible linkers (15–30 amino acids)^26^. Plasmids for protein purification were constructed using pE-SUMO as the backbone. All DNA fragments were amplified by PCR using high-fidelity polymerase with the primers listed in the table, purified by gel extraction, and then assembled into the restriction enzyme-linearized target vectors by commercial homologous recombination cloning kit. All constructs were verified by sequencing, de-methylated in *E. coli* JM110, and then transformed into haloarchaea by PEG-mediated spheroplast transformation^24^.

### *In vivo* Crosslinking and IP-MS to screen for division proteins

*H. volcanii* H26 strain expressing GFP-fused CdpB1 under the control of the *P_tna_* promoter was used as the test group, while a strain expressing GFP alone served as the negative control. Single colonies were inoculated into 5 mL of Hv-YPCTE medium and cultured at 45 °C for 48 h. The cultures were then diluted 1:100 (v/v) into 80 mL of fresh Hv-YPCTE medium supplemented with 1 mM tryptophan (Trp) (three biological replicates per group) and grown at 45 °C for approximately 12 h until an OD_600_∼0.5 was reached. Cells were harvested by centrifugation at 10,000 rpm for 10 min at 4 °C, and the pellets washed with 16 mL of high-salt PBS buffer (HSPB; PBS containing 144 g/L NaCl. Crosslinking was performed by addition of 640 μL of 25 mM DSP (3,3’-Dithiobis(succinimidyl propionate)) dissolved in DMSO, followed by incubation on ice for 2 h. Subsequently, the reaction was quenched by addition of 2 mL of 1 M Tris-HCl (pH 8.2). The cultures were gently mixed by inversion, and incubated at room temperature for 15 min. Cells were then washed once with 16 mL of HSPB. The cell pellet was then resuspended in 1 mL of HKPBT buffer (HKPB with 0.5% (v/v) Tween-20; HKPB: PBS containing 194 g/L KCl) and lysed by sonication. The lysate was centrifuged to remove the cell debris, and 200 μL of the supernatant was taken as the input sample for Western blot analysis to assess crosslinking efficiency.

For each IP reaction, 800 μL of the supernatant was mixed with GFP antibody at a 1:200 dilution and incubated at 4 °C for 8 h with gentle agitation. Meanwhile, 80 μL of magnetic beads were pre-washed twice with 800 μL of PBST (PBS with 0.5% Tween-20) and twice with HKPBT. The supernatant-antibody mixture was then combined with the pre-washed beads and incubated overnight at 4 °C. After incubation, the supernatant was collected, and the beads were washed three times with HKPBT and three times with HKPB (without Tween-20). One quarter of the beads were reserved for examination of precipitation efficiency, while the remaining three quarters were briefly dried by aspiration and stored at –80 °C for subsequent mass spectrometry analysis.

For Western blot analysis, the reserved beads (one quarter) were eluted with 30 μL of 1× SDS-PAGE loading buffer. A 10 μL aliquot was kept as the non-reduced sample. The remaining 20 μL was mixed with 1 μL of 1 M DTT (final concentration ∼50 mM), heated at 95 °C for 10 min to reduce and reverse crosslinks, and used as the reduced sample. Input, post-IP supernatant, non-reduced eluate, and reduced eluate from each replicate were analyzed by immunoblotting to verify successful immunoprecipitation and un-crosslinked immunocomplexes. If no issues were detected in the efficiency test, the frozen beads were submitted to the SpecAlly Life Technology Co. (Wuhan, China) for LC-MS/MS analysis.

Proteins isolated with the bait (CdpB1-GFP) with a fold change > 4 over the control (GFP) were considered as interactors of the bait. For each candidate interactor gene, N-terminal and C-terminal GFP fusions were constructed. Their localization was assessed in H26 cells, and proteins with clear midcell localization were prioritized for further studies.

### Microscopic observation

Before microscopic observation, all strains were transferred to Hv-Cab medium and cultured overnight twice at 45 °C with shaking at 200 rpm. For fluorescent fusions under the control of the *P_tna_* promoter, 0.2 mM Trp was added to the culture during the overnight cultivation. Cells were harvested at mid-exponential phase (OD_600_ ∼0.2-0.4). A 2 μl aliquot was placed on a 1.5% agarose pad prepared with 18% BSW (Hv-Ca without casamino acids and CaCl₂) and covered with a coverslip.

Images were acquired using an Olympus BX53 upright microscope equipped with a Retiga R1 camera, a CoolLED pE-4000 light source, and a U Plan XApochromat 100× oil-immersion objective (1.45 numerical aperture). GFP and mCherry were visualized using Chroma filter sets 49002 and 49008, respectively. Image processing was performed with Fiji, Adobe Photoshop 2021 or Illustrator 2023.

### Split-GFP interaction assay

Strains carrying split-GFP plasmids were transferred to Hv-Cab medium and cultured twice overnight at 45 °C with shaking at 200 rpm until the OD_600_ reached approximately 0.5. Subsequently, 0.4 mM Trp was added for induction, and the cultures were further incubated overnight at 30 °C with shaking at 200 rpm. Fluorescence was observed by microscopy as described above. For quantitative measurement, cultures were adjusted to OD_600_ = 1, and 200 μl of each sample was transferred to a 96-well plate. Fluorescence intensity (excitation 489 nm, emission 514 nm) was measured using a Varioskan LUX plate reader. Two biological replicates and three technical replicates were performed for each condition; significance was evaluated by two-tailed Student’s *t*-test.

### 2D-SIM fluorescence microscopy

Strains for 2D-SIM fluorescence microscopy were transferred to Hv-Cab medium and cultured twice overnight at 45 °C with shaking at 200 rpm. Subsequently, 1 mM Trp was added for induction, and the cultures were further incubated overnight at 30 °C with shaking at 200 rpm. Cells were harvested at mid-exponential phase (OD_600_ ∼0.2-0.4). A 3.5 μL aliquot was placed under a 1.5% agarose pad prepared with 18% BSW in the glass bottom dish (Cellvis). The cells were then imaged with a HIS-SIM (High Intelligent and Sensitive Structured-Illumination Microscopy) microscope (CSR Biotech) in 2D-SIM mode using 488 nm laser excitation with a ×100/1.47 NA oil-immersion objective lens. Raw images were processed by Wiener reconstruction followed by Sparse reconstruction to obtain super-resolution images. Super-resolution images were processed using Fiji software.

### Western blotting

Prepared protein samples were separated on 12% polyacrylamide gels and were transferred to nitrocellulose (NC) membranes (Pall). Membranes were blocked with 5% skim milk for 1 h at room temperature, then incubated overnight at 4 °C with primary antibody (1:10,000 dilution). After five washes with TBST, membranes were incubated with HRP-conjugated secondary antibody (1:10,000 dilution) for 1 h at room temperature. Signals were developed using an ultra-sensitive chemiluminescent substrate and imaged with a ChemiDoc system.

### Co-immunoprecipitation

Cells expressing the indicated tagged proteins were grown in 40 mL of Hv-YPC medium at 45 °C with shaking at 200 rpm until OD_600_ reached approximately 1.0. The cultures were then transferred to 30 °C with shaking at 200 rpm and incubated overnight with 1 mM Trp to induce expression. Cells were harvested by centrifugation at 10,000 rpm for 10 min at 4 °C, and the pellet was resuspended in 2 ml of HKPBT containing protease inhibitor cocktail and lysed by sonication. The lysate was centrifuged at 10,000 rpm for 10 min at 4 °C. For the input sample, 200 μL of the supernatant was mixed with 50 μL of 5× SDS-PAGE loading buffer and boiled at 95 °C for 10 min. Meanwhile, 400 μL of the supernatant was incubated with magnetic beads coated with antibodies against GFP, Flag, or His (as indicated) overnight at 4 °C. The beads were washed three times with HKPBT, and the immunocomplexes were eluted by boiling in 1× SDS-PAGE loading buffer. The eluted samples, together with the input samples, were analyzed by Western blot as described above.

### Protein expression level analysis

*H. volcanii* H26 strains carrying plasmids expressing GFP fusions of wild-type CdpC or its mutants under the control of the *P_tna_* promoter were inoculated into 40 mL of Hv-YPCTE medium and grown at 45 °C with shaking at 200 rpm for 16 h. Expression was then induced by addition of 1 mM Trp, and the cultures were further incubated overnight at 30 °C with shaking at 200 rpm. After measuring the OD_600_, the cell cultures were normalized across samples. An equal volume of each culture was harvested by centrifugation at 10,000 rpm for 10 min at 4 °C. The pellets were washed once with 10 mL of HSPB and then resuspended in 2 mL of HKPB supplemented with protease inhibitor cocktail. Cells were lysed by sonication, and the lysates were centrifugated to remove cell debris. For each sample, 200 μL of the supernatant was mixed with 50 μL of 5× SDS-PAGE loading buffer and heated at 95 °C for 10 min to prepare the samples. Western blot analysis was then performed as described above.

### Protein purification

Proteins were expressed heterologously in *E. coli* BL21 (DE3) carrying plasmids pZS246 (H-FtsZ1), pZS321 (H-SUMO-FtsZ1), or pZWC511 (H-SUMO-CdpC). Overnight culture harboring plasmid expressing the respective protein was diluted 1:100 into 300 ml of fresh LB, then grown at 37 °C until OD_600_ reached 0.4. Protein expression was induced by addition of IPTG to a final concentration of 1 mM, followed by further incubation for 3 h at 37 °C. Following centrifugation, harvested cells were resuspended in 20 ml of high-salt lysis buffer (25 mM Tris-HCl (pH 7.5), 2.6 M KCl, 5% glycerol, 10 mM imidazole, 0.1 mM DTT) and then disrupted by sonication. The lysates were centrifuged at 18,000 g for 10 min at 4 °C. The supernatants were then applied onto a pre-equilibrated Ni-NTA agarose column. Following a single wash with high-salt wash buffer (25 mM Tris-HCl (pH 7.5), 2.6 M KCl, 5% glycerol, 20 mM imidazole, 0.1 mM DTT), the retained proteins were eluted using elution bufferr (25 mM Tris-HCl pH 7.5, 2.5 M KCl, 5% glycerol, 250 mM imidazole, 0.1 mM DTT). Peak fractions identified by SDS-PAGE were pooled, dialyzed against storage buffer (25 mM Tris-HCl pH 7.5, 2.5 M KCl, 5% glycerol, 0.1 mM DTT), aliquoted, and frozen at -80 °C.

The H-SUMO tag was cleaved from CdpC or FtsZ1 by purified 6×His-tagged SUMO protease (Ulp1) in the protein storage buffer with 200 mM KCl at 30 °C for 1 h. he cleavage mixture was then passed through fresh Ni-NTA to retain the released His-SUMO tag and the His-tagged protease, while the untagged target protein was recovered in the flow-through. After dialysis against storage buffer and concentrated, if needed. The concentration of purified proteins was determined and the proteins were stored at -80 °C. His-tagged CdpC_CTD_ was synthesized commercially by GenScript Biotech Corporation (China).

### Biolayer interferometry assay

BLI measurements were carried out on an Octet-Red96 system at 30 °C. To assess the interaction between FtsZ1 and either CdpC or its C-terminal domain (CdpC_CTD_), Ni-NTA biosensors were first pre-equilibrated in 1×Pol buffer (12.5 mM HEPES-NaOH [pH 6.8], 2.6 M KCl, 2.5 mM MgCl2, 0.02% Tween-20). Then, 6×His-tagged FtsZ1 or 6×His-tagged CdpC_CTD_ (each at 0.5 μM) was loaded onto the biosensor tips for 3 min using 200 μL of the same buffer supplemented with GTP to a final concentration of 1 mM, followed by a 60-second wash to eliminate unbound or weakly attached protein. The association of untagged CdpC with the immobilized His-FtsZ1 was recorded over 4 min under shaking at 1,000 rpm, and dissociation was then allowed to proceed for 3 min in buffer without any protein. To examine the binding of FtsZ1 to 6×His-CdpC_CTD_, 6×His-CdpC_CTD_ was immobilized onto the biosensors as described above, and the binding of untagged FtsZ1 was followed for 5 min, after which dissociation was monitored for 3 min. Data analysis and curve fitting were performed using the ForteBio Data Analysis Software and GraphPad Prism 10.

### Statistical analysis

GraphPad Prism 10 was used for all statistical tests. Two-group comparisons were performed using two-tailed, unpaired Student’s *t*-test. A P-value < 0.05 was considered significant. All experiments were repeated at least three independent times with similar results. Data are presented as mean ± standard deviation (SD). Sample sizes (n) are indicated in the figure legends.

### Homolog search

An iterative HMM search approach as described previously^31,37^ was used to find sequence homologs of CdpC across archaea, with an updated database comprising GTDB r226^38^ and additional Asgard archaeal dataset as described in ref^31^. Briefly, HvCdpC was used as seed to search for homologs via Diamond v2.1.13.167. Then all the hits were aligned to confirm the presence of critical domains, and when uncertainty arose, AlphaFold structural predictions were used for an additional confirmation. Sequences were clustered using mmSeqs2 v13.45111^39^ with sequence identity threshold of 0.4, then aligned by MAFFT v7.505^40^ (“auto” mode), pruned by TrimAl ^41^ v1.4 (“gappyout”), and a preliminary tree was constructed using FastTree v2.1.11 (default parameter)^42^. Afterwards, tree leaves were reduced to 100 by Treemmer v0.3. The sequences were aligned and pruned again using MAFFT “L-INS-i” mode, manually examined to remove truncated or poorly aligned sequences, and then used to build the HMM profile using HMMer v3.3.2 function “hmmbuild” (http://hmmer.org/). We then used the profile to search for homologs within the database above using HMMer function “hmmsearch”. The above processes were iterated 3 times to obtain the final profile and search results.

To search for remote homologs of Halobacterial CdpC across archaea, we first examined the individual sequences around threshold and carried out AF3 structural prediction and found that they indeed pertained one or more zinc ribbon domains. We then used Diamond search to find homologs of these proteins. A combined HMM profiles based on a combined set of these homologs and Halobacterial CdpC homologs were used to search against the above archaeal proteome database. Representatives from each archaeal phylum were selected for AF3 prediction.

### Phylogenetic and alignment fraction analysis

All sequences of Halobacterial CdpC homologs were aligned using MAFFT “L-INS-i” mode. The sequence identity of the MSA was calculated and visualized using a custom matlab script. The MSA was then trimmed using TrimAl option “gappyout”, and then analyzed using IQ-TREE v2.2.2.2 option “-m LG +F+G4 -B 2000”. The tree was visualized on the itol website^43^.

### Protein structural prediction

Protein structures were predicted. All candidate protein sequences identified from the HMM-based searches were modeled with the default parameter set using the Protenix online server (Protenix, ByteDance Inc.)^44^. Both the MSA-enabled mode and the template-enabled mode were activated to incorporate evolutionary information and homologous structural features during prediction. For each query, the corresponding amino acid sequence was submitted together with Zn²⁺ ions specified as ligands to guide the modeling of zinc-binding motifs. The resulting structural models were visualized and inspected directly within the Protenix platform.

## Acknowledgement

We thank members of the Du lab, Chen lab and Wu lab for advice and helpful discussions to carry out this study. This study was supported by National Natural Science Foundation of China (grant 32270049 and 32070032), and the Young Top-notch Talent Cultivation Program of China to S.D.; National Natural Science Foundation of China (grant 323B2001) to S.Z.; Research in the Wu lab was supported by National Natural Science Foundation of China (grant 32570019 and 32370003).

## Data of Availability

Data generated and analyzed during this study are presented in the paper or in the Supplementary Information and Datasets. Plasmids and strains that support the findings of this study are available from the corresponding authors.

## Extended Data Figure Legends

**Extended Data Fig 1.**
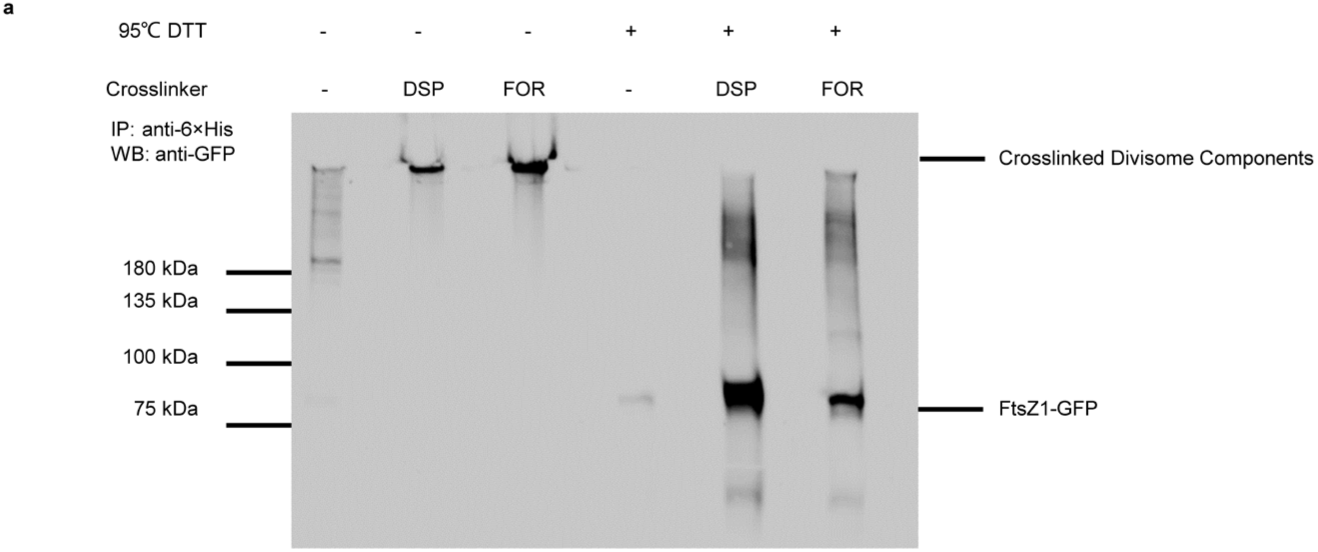
*In vivo* crosslinking facilitates the isolation of indirect interaction partners of a division protein. Cultures of *H. volcanii* expressing FtsZ1-GFP and CdpB1-6×His were treated with or without the crosslinker formaldehyde (FOR) or DSP. Cells were lysed by sonication; supernatants were incubated with anti-His antibodies coated magnetic beads. Immunocomplexes were treated with DTT (final concentration 50 mM, 95°C, 10 min) or directly eluted with SDS-PAGE loading buffer and then analyzed by immunoblot.

**Extended Data Fig. 2.**
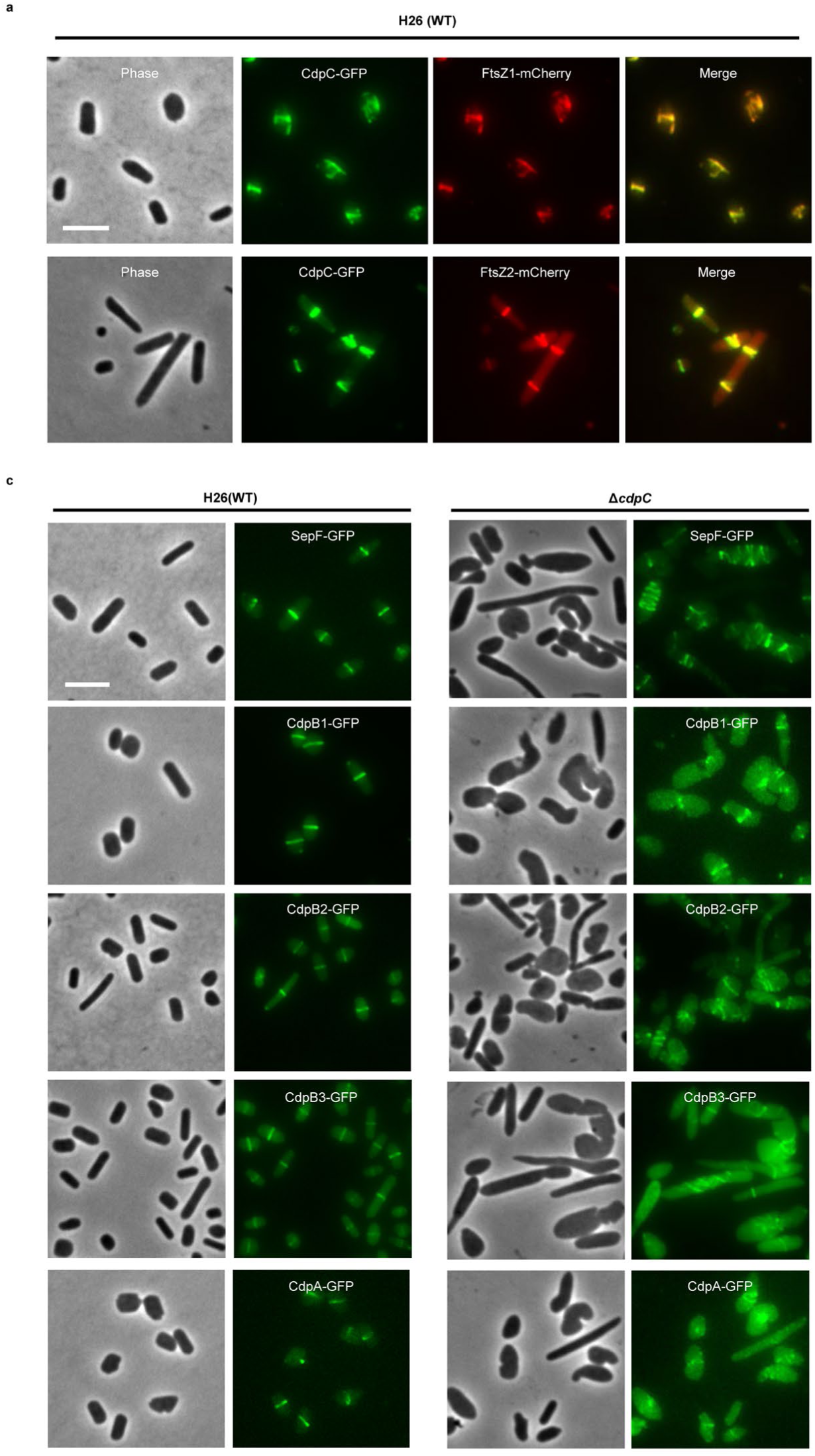
Co-localization of CdpC with FtsZ1/2 and the impact of cdpC deletion on localization of known division proteins. a. Representative images of the co-localization of CdpC-GFP with FtsZ1-mCherry or FtsZ2-mCherry in H26(WT) cells. Cells were cultured in Hv-Cab medium at 45 °C, CdpC-GFP and FtsZ1-mCherry or FtsZ2-mCherry were co-expressed from the same plasmid. Expression of FtsZ1-mCherry or FtsZ2-mCherry was driven by their respective native promoters on the plasmid, while CdpC-GFP was under the control of the *P_tna_* promoter and induced with 0.2 mM tryptophan (Trp) overnight. Scale bar, 5 μm. b. Deletion of *cdpC* affects localization of known cell division proteins. Cells were cultured in Hv-Cab medium at 45 °C, SepF-GFP and CdpB1-GFP were expressed from their native promoters on plasmids, whereas CdpB2-GFP, CdpB3-GFP, and CdpA-GFP were driven by the *P_tna_* promoter on plasmids and induced with 0.2 mM Trp overnight. Scale bar, 5 μm.

**Extended Data Fig. 3.**
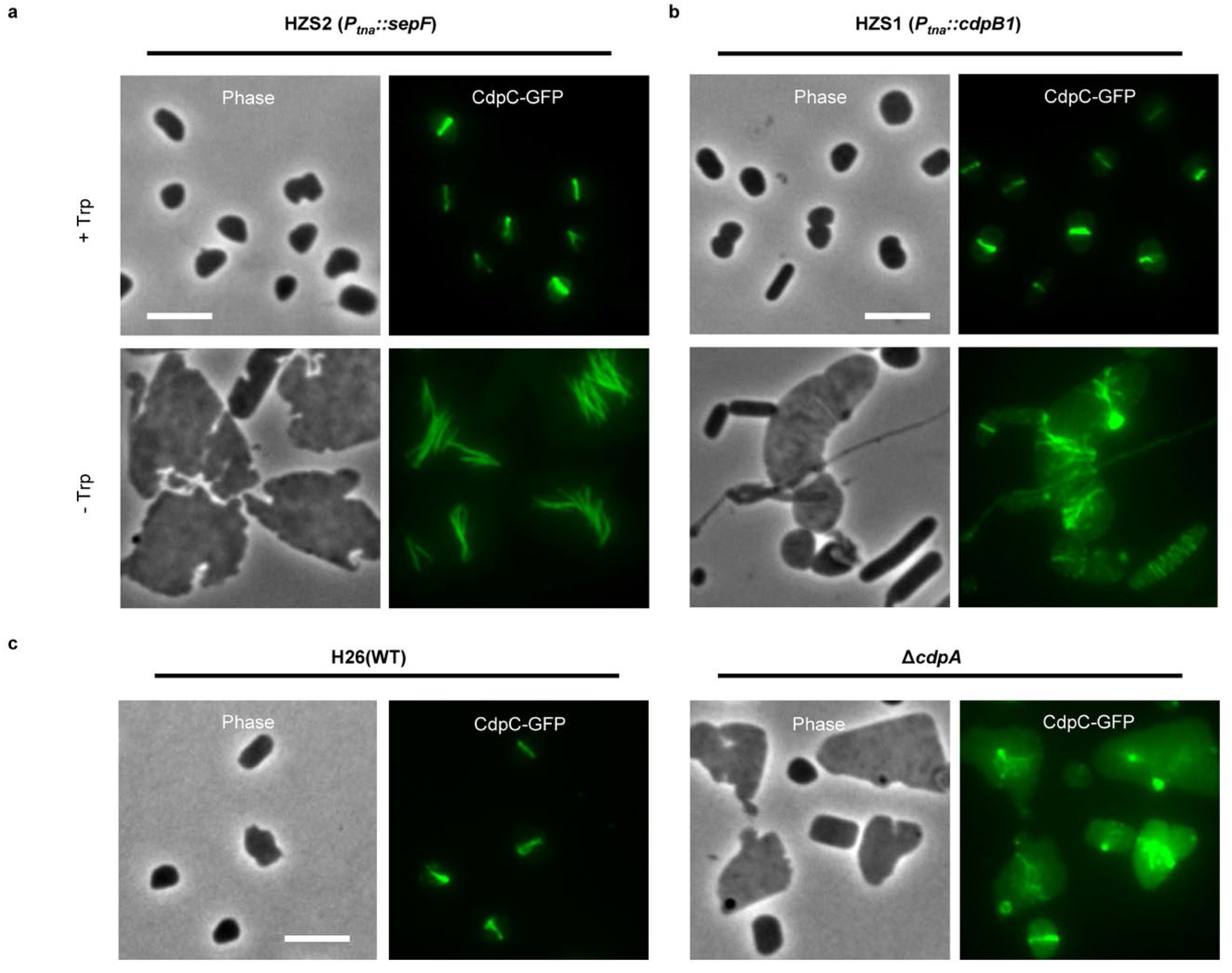
CdpC does not depend on SepF, CdpA or CdpB1 for localization. a-b. Representative images of the localization of CdpC-GFP in the presence or absence of SepF (a) or CdpB1 (b). Cells were cultured in Hv-Cab medium at 45 °C, expression of CdpC-GFP was driven by its native promoter on plasmid. Depletion of SepF or CdpB1 was achieved by removal of tryptophan (Trp) from the cultures. c. Representative images of the localization of CdpC-GFP in wild type and *ΔcdpA* cells. Cells were cultured in Hv-Cab medium at 45 °C, expression of CdpC-GFP was driven by its native promoter on plasmid. Scale bars, 5 μm.

**Extended Data Fig. 4.**
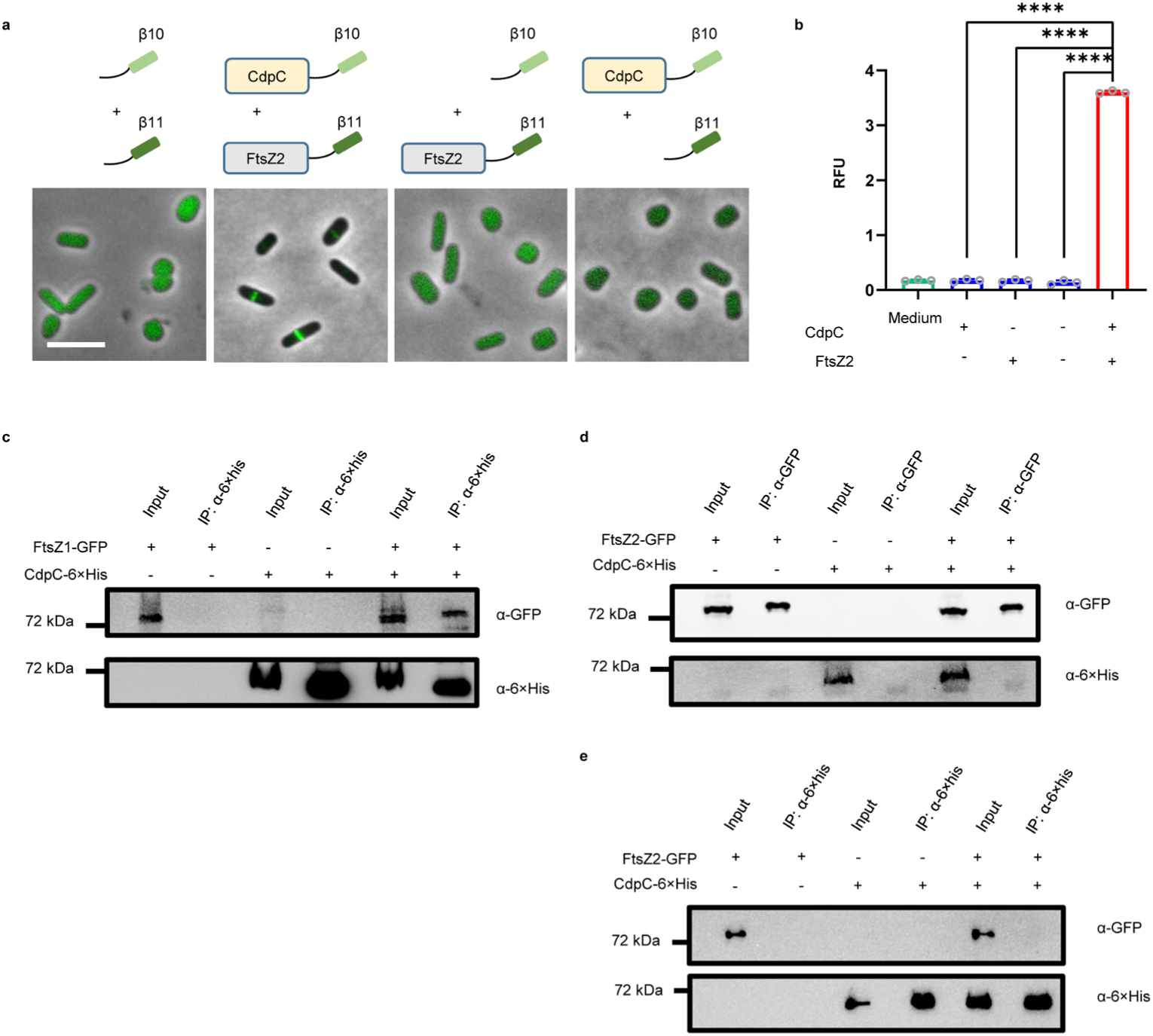
CdpC interacts with FtsZ1 but not with FtsZ2. a. Representative images showing the fluorescence and localization of sfGFP of Split-FP assay. Scale bar, 5 μm. b. Quantitation of the interaction signal between CdpC and FtsZ2 by Split-FP assay in H26 (WT) cells. RFU, relative fluorescence unit. Data are presented as mean values ± s.d. Significance in each group was tested by two-sided *t*-test. \*\*\*\**P* <0.0001, *n*=3. c. Co-IP experiment showing that FtsZ1-GFP co-immunoprecipitated with CdpC-6×his *in vivo*. Cultures of *H. volcanii* expressing the indicated proteins were lysed by sonication; supernatants were incubated with anti-6×his antibodies coated magnetic beads. Immunocomplexes were eluted with boiling SDS–PAGE loading buffer and then analyzed by immunoblot. d-e. Co-IP experiments showing that CdpC does not interact with FtsZ2 *in vivo*. The assay was carried out as in c with anti-GFP or anti-His antibodies coated magnetic beads.

**Extended Data Fig. 5.**
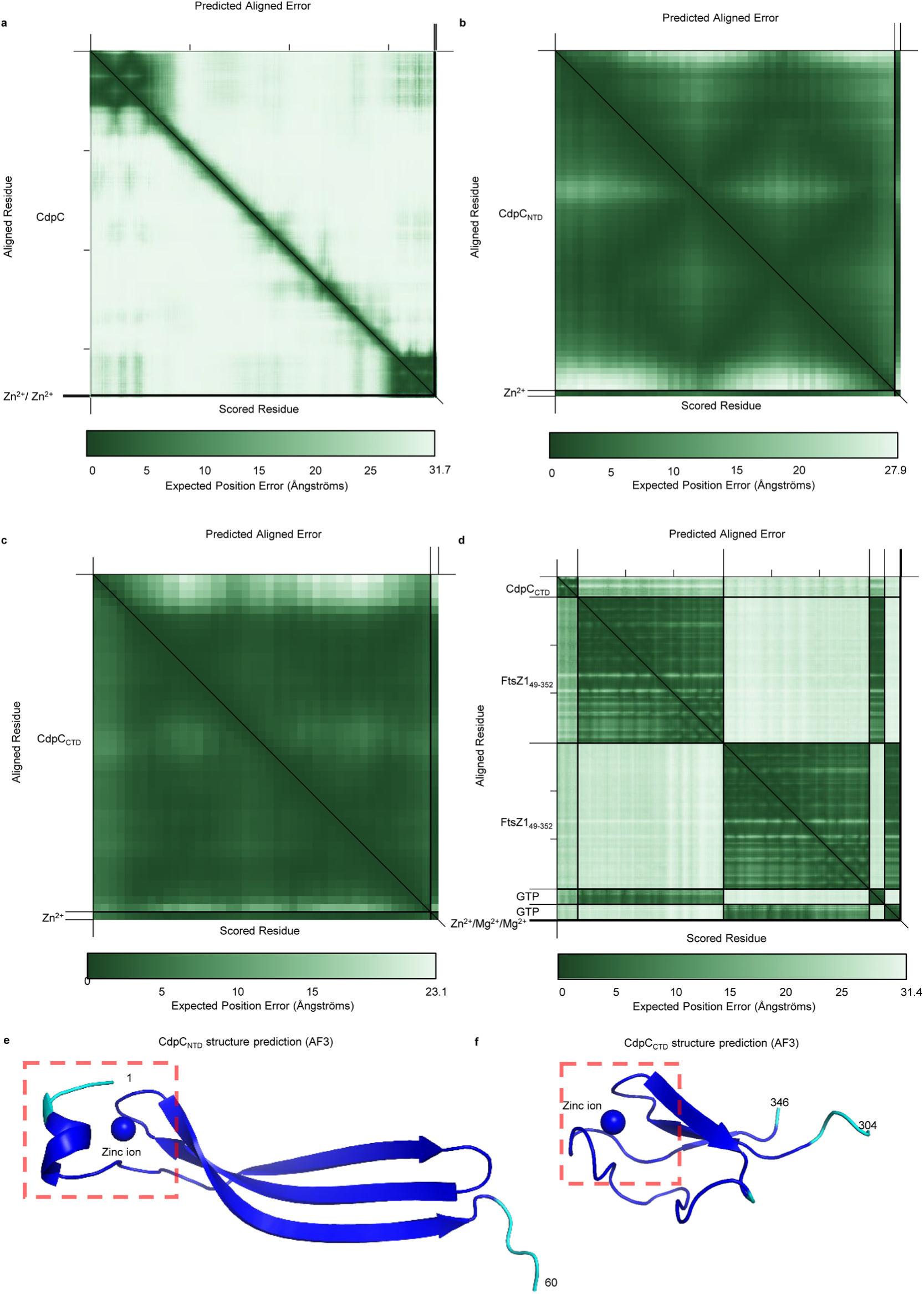
The Alphafold3 structure prediction of full length CdpC, CdpC_NTD_, CdpC_CTD_ and CdpC_CTD_-FtsZ1 complex. a-d. The Alphafold 3 predicted aligned error (%) heatmap plot of full length CdpC, CdpC_NTD_, CdpC_CTD_ and CdpC_CTD_-FtsZ1 complex. The degree of confidence in pairwise interactions between residues was shown. e-f. Alphafold3 structural model of CdpC_NTD_ with one bound zinc ion colored by model confidence (blue - pLDDT > 90%, cyan - 90% > pLDDT > 70%, yellow – 70% > pLDDT > 50%, red - pLDDT <50%), ipTM = 0.91, pTM=0.66. The zinc finger is indicated by a red box. Zinc ion is depicted as a blue ball. f. Alphafold3 structural model of CdpC_CTD_ with one bound zinc ion colored by model confidence (blue - pLDDT > 90%, cyan - 90% > pLDDT > 70%, yellow – 70% > pLDDT > 50%, red - pLDDT <50%), ipTM = 0.86, pTM=0.7. The zinc finger is indicated by a red box. Zinc ion is depicted as a blue ball.

**Extended Data Fig. 6.**
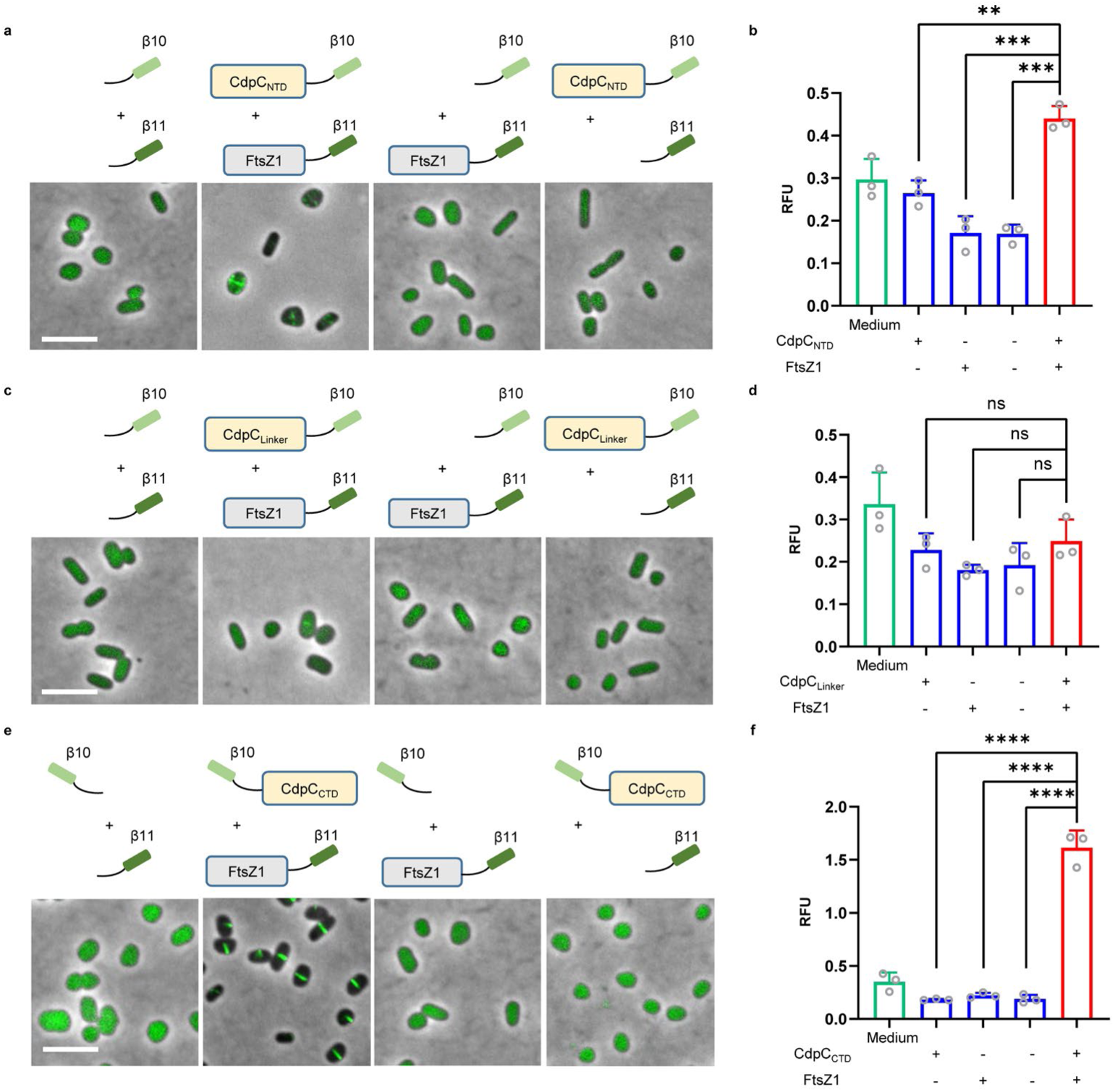
Examination of the interaction between FtsZ1 and the individual domains of CdpC by Split-FP assays. a. Representative images of cells co-expressing tagged CdpC_NTD_ and FtsZ1 in Split-FP assay. b. Quantitation of the interaction signal between CdpC_NTD_ and FtsZ1 by Split-FP assay in H26 (WT). RFU, relative fluorescence unit. Data are presented as mean values ± s.d. Significance in each group was tested by two-sided *t*-test. ** *P* <0.01, *** *P* <0.001, *n*=3. c. Representative images of co-expressing tagged CdpC_Linker_ and FtsZ1 in Split-FP assay. d. Quantitation of the interaction signal between CdpC_Linker_ and FtsZ1 by Split-FP assay in H26 (WT). RFU, relative fluorescence unit. Data are presented as mean values ± s.d. Significance in each group was tested by two-sided *t*-test. ns *P* >0.05, *n*=3. e. Representative images of co-expressing tagged CdpC_CTD_ and FtsZ1 in Split-FP assay. f. Quantitation of the interaction signal between CdpC_CTD_ and FtsZ1 by Split-FP assay in H26 (WT). RFU, relative fluorescence unit. Data are presented as mean values ± s.d. Significance in each group was tested by two-sided *t*-test. \*\*\*\**P* <0.0001, *n*=3. Scale bars, 5 μm.

**Extended Data Fig. 7.**
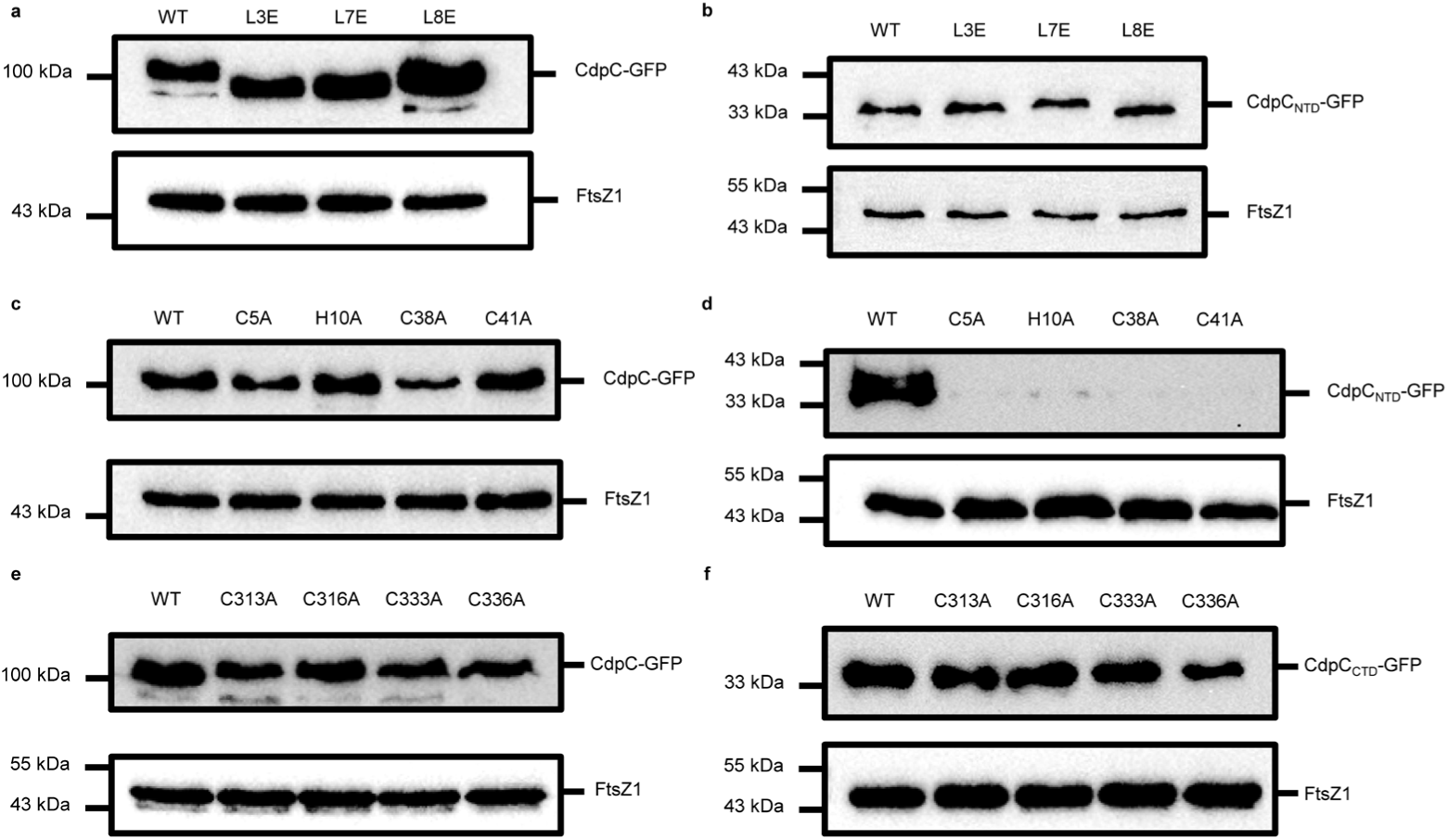
Western blot to access the level of different CdpC variants. a. Western blot analysis of the expression levels of CdpC-GFP variants with mutations in the amphipathic helix. GFP fusions of CdpC and its mutants were expressed from plasmids under the control of the *P_tna_* promoter, induced with 1 mM tryptophan (Trp) overnight at 45 °C. Cultures were normalized to the same OD_600_, then lysed by sonication. The supernatants were mixed with 5× SDS-PAGE loading buffer, boiled for 10 min, and analyzed by immunoblotting. FtsZ1 serves as the loading control. b. Western blot analysis of the expression levels of CdpC_NTD_-GFP variants with mutations in the amphipathic helix. The assay was carried out as in a. FtsZ1 serves as the loading control. c. Western blot analysis of the expression levels of CdpC-GFP variants with mutations in the zinc-finger motif in the NTD. The assay was carried out as in a. FtsZ1 serves as the loading control. d. Western blot analysis of the expression levels of CdpC_NTD_-GFP variants with mutations in the zinc-finger motif in the NTD. The assay was carried out as in a. FtsZ1 serves as the loading control. e. Western blot analysis of the expression levels of CdpC-GFP variants with mutations in the zinc-finger motif in the CTD. The assay was carried out as in a. FtsZ1 serves as the loading control. f. Western blot analysis of the expression levels of CdpC_CTD_-GFP variants with mutations in the zinc-finger motif in the CTD. The assay was carried out as in a. FtsZ1 serves as the loading control.

**Extended Data Fig. 8.**
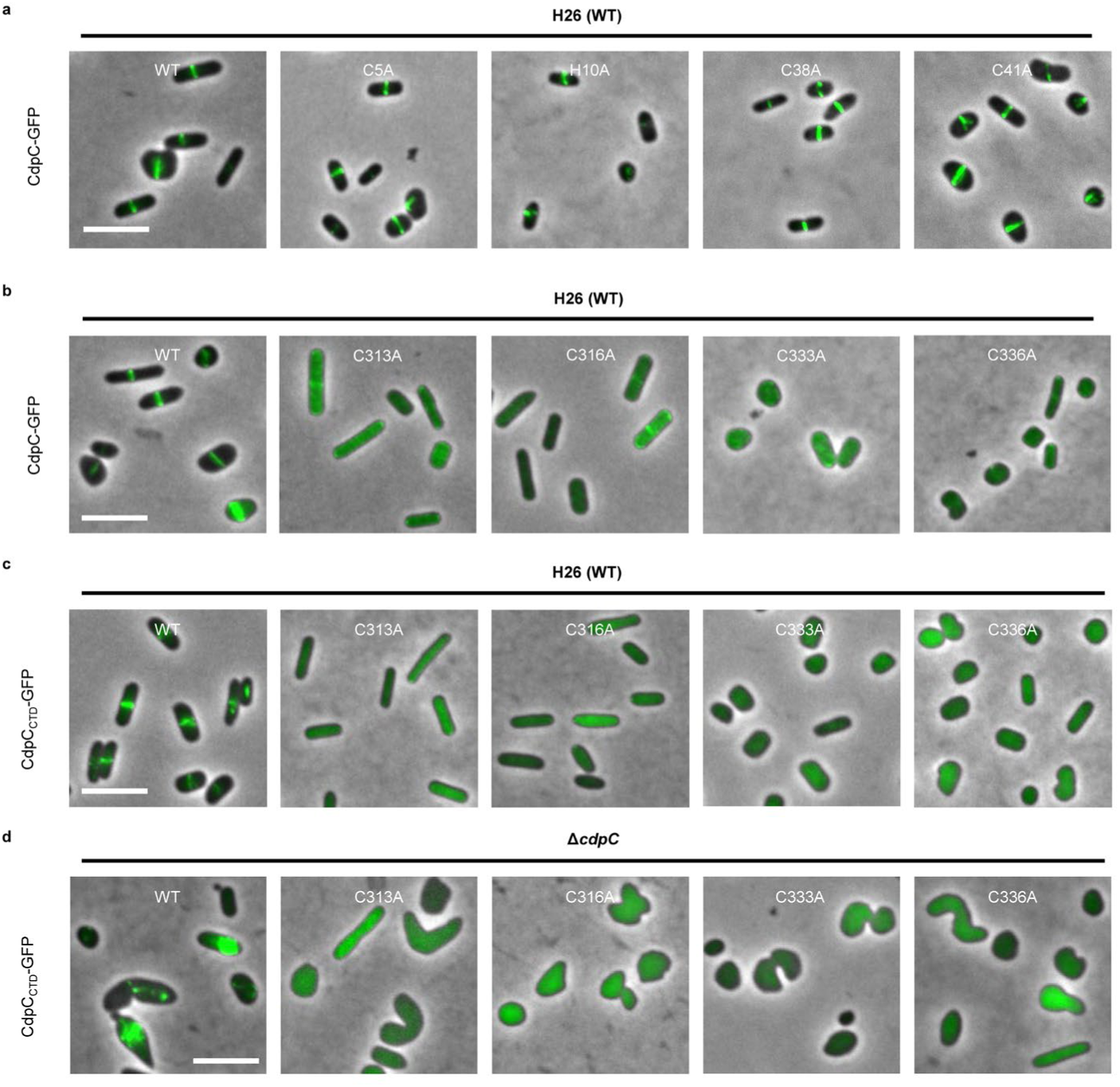
Disruption of the zinc finger in the CTD abolishes the midcell localization of CdpC. a. Representative images showing that mutations in the zinc finger in the NTD do not disrupt the midcell localization of CdpC-GFP in H26 (WT) cells. Cells were cultured in Hv-Cab medium at 45 °C, GFP fusion of wild type CdpC and its mutants were expressed from plasmids under the control of the *P_tna_* promoter, induced with 1 mM tryptophan (Trp) overnight. b. Representative images showing that mutations in the zinc finger in the CTD disrupt the midcell localization of CdpC-GFP in H26 (WT) cells. The experiment was carried out as in a. c. Representative images showing that mutations in the zinc finger disrupt the midcell localization of CdpC_CTD_-GFP in H26 (WT) cells. The experiment was carried out as in a. d. Representative images showing that mutations in the zinc finger disrupt the filamentous localization of CdpC_CTD_-GFP in *Δ*cdpC cells. The experiment was carried out as in a. Scale bars, 5 μm.

**Extended Data Fig. 9.**
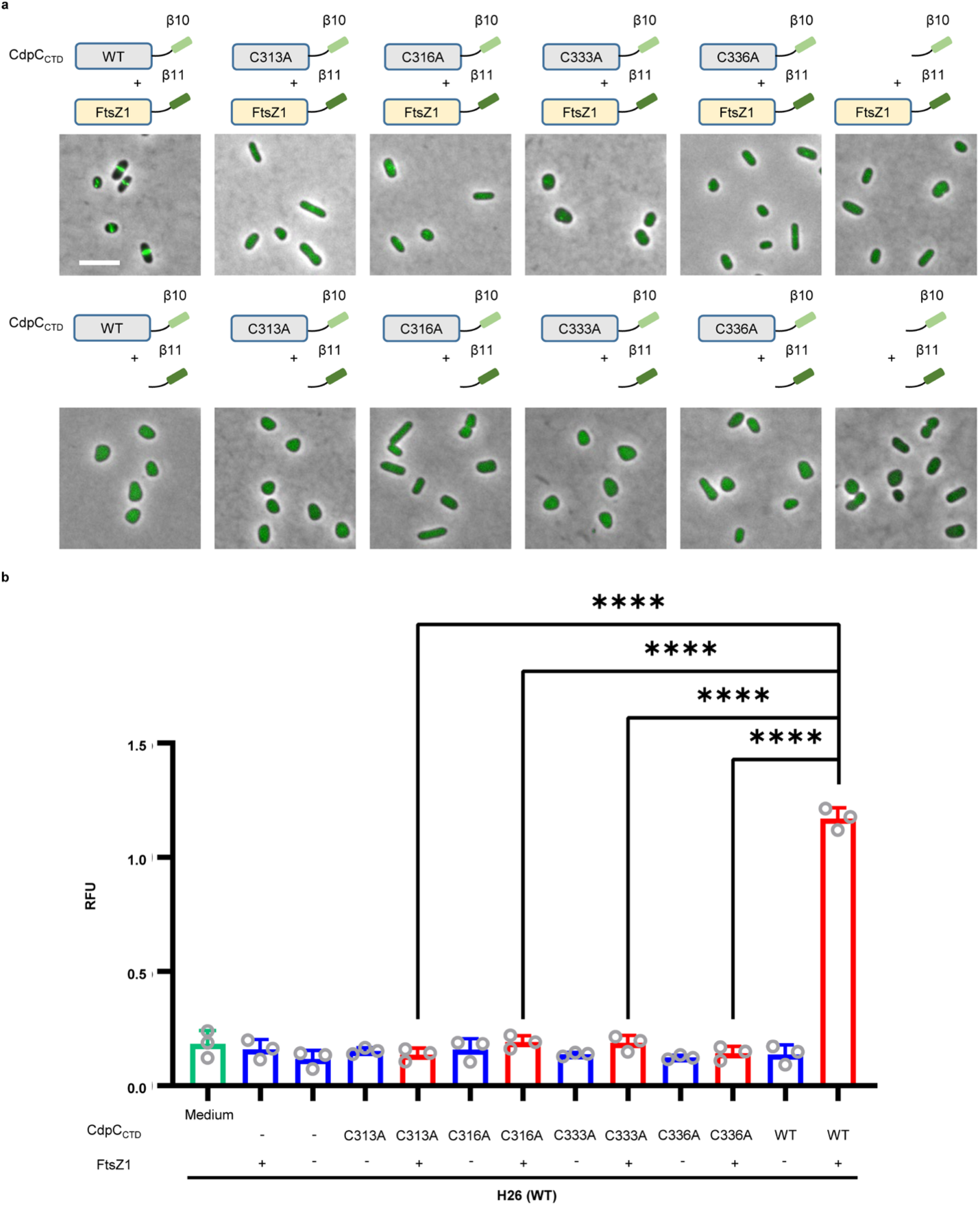
Disruption of the zinc finger abolishes the interaction between CdpC_CTD_ and FtsZ1. a. Representative images of Split-FP assay to access the interaction between the mutants of CdpC_CTD_ and FtsZ1 *in vivo*. Scale bar, 5 μm. b. Quantitation of the interaction signal between the mutants of CdpC_CTD_ and FtsZ1 by Split-FP assay. RFU, relative fluorescence unit. Data are presented as mean values ± s.d. Significance in each group was tested by two-sided *t*-test. \*\*\*\**P* <0.0001, *n*=3.

**Extended Data Fig. 10.**
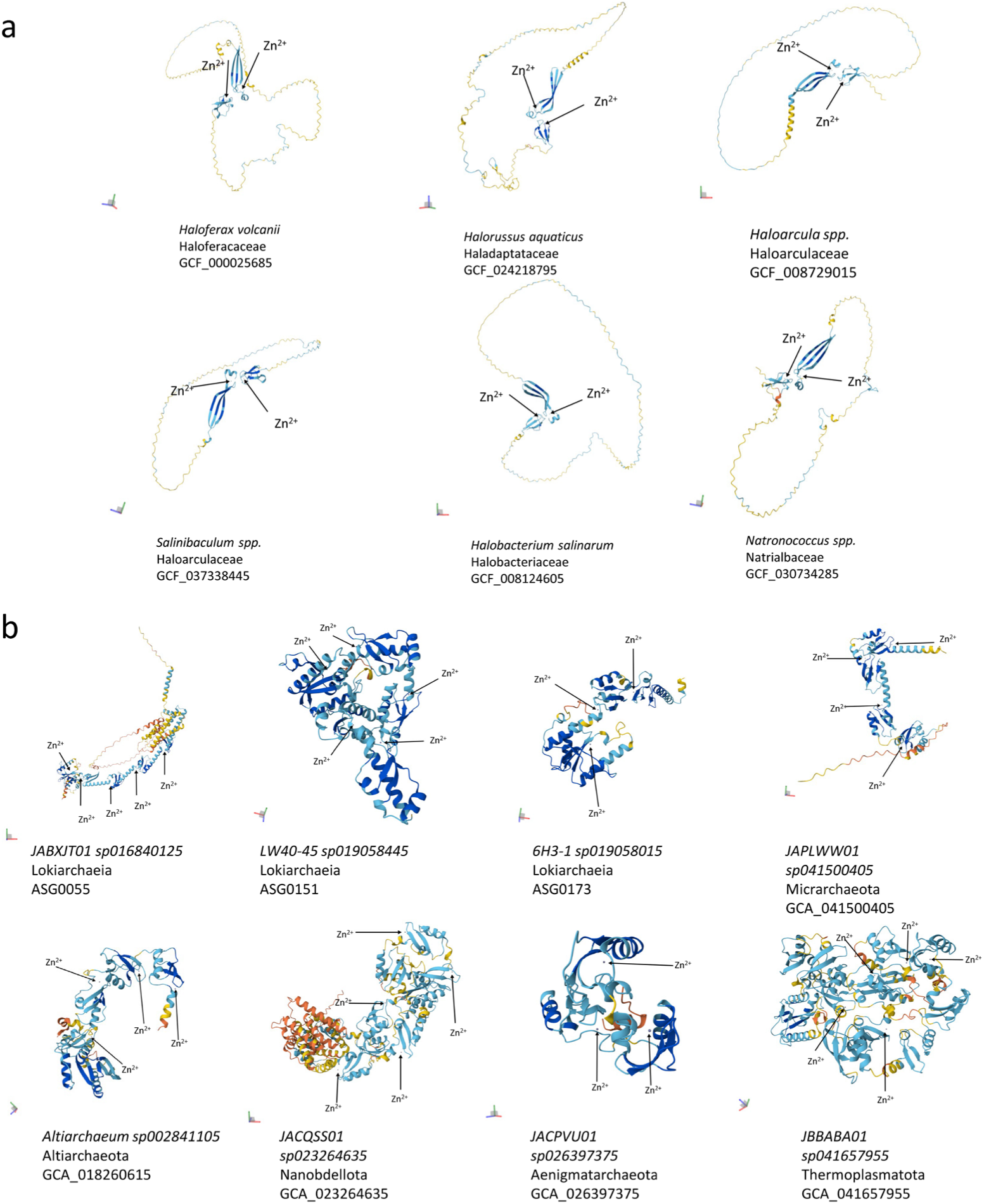
Structural features of CdpC homologs in archaea. a. AF3 structural prediction of 6 CdpC homologs from 6 different families of *Halobacteria*. Predicted locations of the zinc ion are indicated with arrow. b. AF3 structural prediction of 8 distant homologs of CdpC from 6 different phyla. Predicted locations of the zinc ion are indicated with arrow.

